# DPF3a couples H3K36me2-dependent chromatin remodeling to genome architecture during myogenesis

**DOI:** 10.64898/2026.08.25.746968

**Authors:** Ming Yu, Jingdong Xue, Qi Zhang, Yixuan Pan, Min Hao, Mingqian Hu, Meiyang Liu, Yuqian Feng, Yanhua Yao, Mengyuan Peng, Jun Wu, Yupeng Chen, Ping Hu, Yimin Lao, Bing Li

## Abstract

Chromatin remodelers are generally thought to regulate gene expression through nucleosome mobilization and modulation of local chromatin accessibility. Whether these complexes can instead control transcription through higher-order chromatin organization remains poorly understood. Here we identify the BAF subunit DPF3a as an essential regulator of skeletal muscle regeneration and myogenic differentiation. DPF3a functions through association with the H3K36me2/3 reader HRP2 and preferentially localizes to a distinctive chromatin state characterized by focal depressions within broad H3K36me2 domains. Biochemical reconstitution demonstrates that H3K36 methylation directly enhances remodeling activity of the DPF3a-containing cBAF complex. Unexpectedly, despite profound transcriptional defects, loss of DPF3a produces minimal changes in local chromatin accessibility. Instead, DPF3a is required for long-range chromatin looping associated with activation of myogenic genes. Together, our findings uncover a non-canonical mechanism whereby a chromatin remodeler regulates transcription primarily through three-dimensional genome organization rather than local accessibility control, and establish histone modification-guided chromatin remodeling as a key principle in gene regulation.

**Key Points:**

- DPF3a remodels chromatin through genome looping rather than promoter accessibility
- H3K36me2/3-directed cBAF remodeling governs myogenic gene regulation
- DPF3a controls skeletal muscle regeneration through chromatin architecture

## Introduction

The mammalian SWI/SNF (switch/sucrose non-fermentable) chromatin remodeling complexes, also known as BAF (BRG1/BRM-associated factor) complexes, are ATP-dependent molecular machines that reposition nucleosomes to regulate chromatin organization and gene expression ^1–5^. Through ATP hydrolysis-driven remodeling, BAF complexes generate chromatin environments permissive for transcription factor binding and RNA polymerase II (RNA Pol II) engagement^6–10^. Mammalian BAF complexes comprise three principal assemblies—canonical BAF (cBAF), polybromo-associated BAF (PBAF), and non-canonical BAF (ncBAF)—that share a conserved ATPase core while differing in auxiliary subunit composition. Recent structural and biochemical studies have substantially advanced our understanding of BAF complex assembly, modular organization, and nucleosome engagement^11–19^. Despite these advances, current models largely conceptualize BAF complexes as regulators of local chromatin accessibility through nucleosome mobilization.

Emerging evidence, however, suggests that BAF complexes may also participate in higher-order genome organization. During erythroid differentiation, the ATPase BRG1 was shown to facilitate chromatin looping at the globin locus^20–22^, raising the possibility that BAF-mediated remodeling contributes to long-range chromatin interactions. More recently, nucleosome-resolution chromatin conformation profiling identified several BAF subunits, including DPF2, ARID1B, and SMARCE1, as strong predictors of promoter-centered chromatin contacts^23^. These observations collectively point toward a broader architectural role for BAF complexes beyond canonical accessibility regulation. Nevertheless, the mechanistic relationship between ATP-dependent chromatin remodeling and enhancer-promoter looping remains poorly defined.

Among BAF subunits, DPF3a represents a particularly intriguing and tissue-restricted component of the cBAF complex. Unlike its paralogs DPF1, DPF2, and DPF3b, DPF3a contains a truncated plant homeodomain (PHD) together with a unique C-terminal region^24,25^. DPF3a expression is enriched in skeletal muscle, heart, testis, and ovary (Supplementary Fig. S1a), suggesting specialized lineage-specific functions. Previous studies in the immortalized C2C12 myoblast cell line demonstrated that DPF3a interacts with hepatoma-derived growth factor-related protein 2 (HRP2), a PWWP-domain protein that recognizes H3K36me2 and H3K36me3, and that this interaction contributes to myotube differentiation^26,27^. However, several fundamental questions remain unresolved. First, prior studies primarily emphasized HRP2 function, leaving the physiological role of DPF3a itself incompletely characterized, particularly in primary myoblasts that more faithfully preserve endogenous muscle stem cell properties. Second, although HRP2 recognizes H3K36me2/3, these histone modifications are broadly distributed across the genome, especially H3K36me2, which extensively occupies intergenic domains^28,29^. How the HRP2-DPF3a axis achieves selective genomic targeting within such pervasive chromatin landscapes therefore remains unclear.

Skeletal myogenesis provides a powerful system for examining the interplay between chromatin remodeling, transcriptional regulation, and three-dimensional genome organization. During both embryonic development and adult muscle regeneration, muscle stem cells undergo highly coordinated transcriptional and epigenetic transitions to establish differentiated myofibers^30,31^. These processes require precise integration of histone modifications, transcription factor networks, and chromatin remodeling activities at regulatory elements^30,32^. Notably, dynamic chromatin looping has emerged as a prominent feature of myogenic differentiation. The fast myosin heavy chain (fMyh) locus undergoes extensive reorganization of its chromatin interaction landscape during the myoblast-to-myotube transition^33–35^, while developmental loci such as the Hoxb cluster similarly exhibit differentiation-associated conformational remodeling^36^. Whether BAF complexes directly contribute to these architectural transitions, and whether tissue-specific subunits such as DPF3a mediate such functions, remains unknown.

Here, we investigated the role of DPF3a in primary murine myoblasts and skeletal muscle regeneration using complementary genetic, genomic, biochemical, and chromatin conformation approaches. We find that DPF3a is indispensable for myogenic differentiation and regenerative muscle formation, acting as a central transcriptional co-regulator of muscle gene expression programs. Mechanistically, DPF3a preferentially localizes to focal depressions embedded within broad H3K36me2-enriched plateaus, a distinctive chromatin configuration associated with reduced chromatin density and active regulatory features. DPF3a function requires association with the H3K36me2/3 reader HRP2, and biochemical reconstitution demonstrates that H3K36 methylation directly promotes remodeling activity of the DPF3a-containing cBAF complex. Unexpectedly, despite substantial transcriptional perturbation, loss of DPF3a produces minimal changes in local chromatin accessibility at its functional target loci. Instead, DPF3a is required for long-range chromatin looping at key myogenic regions, including the fMyh and Hoxb loci. Together, these findings establish a non-canonical mechanism by which a chromatin remodeler regulates transcription predominantly through higher-order genome organization rather than local accessibility modulation, and identify histone modification-guided chromatin remodeling as a central principle in lineage-specific gene control.

## Materials and Methods

### Mice

*Dpf3a* knockout mice (strain NO. T053889) were generated by GemPharmatech Co., Ltd (Nanjing, China). CRISPR/Cas9 technology was used to knock out the target sequence at exon 9 of the *Dpf3*-206 (ENSMUST00000177959.8) on the C57BL/6 genetic background. Mice were genotyped by polymerase chain reaction (PCR), and DPF3a deficiency was confirmed by western blot. All experimental mice were mated and reproduced under specific pathogen-free (SPF) conditions. Animal procedures were approved by the Animal Care and Use Committee of Shanghai Jiao Tong University School of Medicine. Animal experimental protocols were reviewed and approved by the Institutional Animal Care and Use Committee of Shanghai Jiao Tong University School of Medicine (Approval No. JUMC2023-068-A).

### Cardiotoxin (CTX)-induced injury model

Mice aged 12 to 16 weeks were anaesthetized and injected intramuscularly with 50 μL of cardiotoxin (10 μM) into tibialis anterior (TA) muscle. Mice were euthanized by cervical dislocation 7 days post-injection. TA muscles were harvested, embedded in Optimal Cutting Temperature (OCT) compound, snap-frozen in liquid nitrogen, and stored at -80°C. Sections (8 μm) were stained with hematoxylin and eosin. Muscle regeneration was assessed by measuring the cross-sectional area of centrally nucleated myofibers from three mice per genotype using Image J.

### Isolation of muscle stem cells

Muscle stem cells were isolated as previously described^37,38^. Briefly, muscle tissues from 2 to 3 weeks mice were minced into ∼1mm^3^ pieces and digested with a mixture of Dispase II (Roche, 4942078001) and collagenase D (Roche, 11088866001) for 90 min at 37℃. The digest was filtered through a 40-μm cell strainer (Corning Falcon, 352340). Erythrocytes were removed using red blood cell lysis buffer (150 mM NH_4_Cl, 10 mM NaHCO_3_, 1 mM EDTA). The cell suspension was stained with a cocktail of APC anti-mouse CD31 (BioLegend, 102510, 1:100), APC anti-mouse CD45 (BioLegend, 103112, 1:100), FITC Ly-6A/E (Sca-1) (BioLegend, 11-5981-85, 1:100), and Biotin anti-mouse VCAM1 (BioLegend, 105704, 1:100) for 45 min at 4 °C. The cells were then washed with PBS containing 0.2% BSA and stained with PE/Cy7 Streptavidin (BioLegend, 405206, 1:100) for 15 min. After an additional wash, cells were stained with propidium iodide (Sigma, P4170) and sorted by FACS using Aria III (BD Biosciences).

### Cell culture and differentiation

Primary myoblasts were cultured with Ham’s F-10 medium (Gibco, 11550043) supplemented with 20% fetal bovine serum (FBS; Gibco, 10099141C), 2.5 ng/mL bFGF (R&D, 233-FB-025), and 1% penicillin-streptomycin (Gibco, 15140122) on collagen-coated dishes at 37°C in 5% CO2. Differentiation was induced by switching to DMEM (BasalMedia, L110KJ) containing 2% horse serum (Gibco, 16050122) and 1% penicillin-streptomycin. Cells were routinely tested and confirmed negative for mycoplasma contamination.

### Immunofluorescence

Cultured cells were fixed in PBS containing 4% paraformaldehyde (Sigma-Aldrich, 16005-1KG-R) for 15 min, permeabilized in 0.5% Triton X-100 for 15 min at room temperature, and blocked in PBS containing 2% BSA (Sigma-Aldrich, V900933-1KG) for 1h at room temperature. Cells were then incubated with anti-MyHC antibody (Sigma-Aldrich, 05-716-I-100UL, 1:1000) overnight at 4°C. After washing, Alexa 546-conjugated anti-mouse secondary antibody (Invitrogen, A11003, 1:1000) was applied for 1 h at room temperature. Finally, samples were mounted in antifade mountant (Invitrogen, P36930) for imaging.

### Western blot

Cells were detached using Trypsin-EDTA (Gibco, 25200056) and washed with PBS. Cell pellets were resuspended in RIPA lysis buffer (1% Igepal CA-630, 0.5% sodium deoxycholate, 0.1% SDS in PBS) freshly supplemented with a protease inhibitor cocktail (Roche, 04693132001), incubated on ice for 30 min, and centrifuged at 13,000 rpm for 15 min at 4°C. The supernatant was collected. Total protein lysates were resolved by SDS-PAGE, transferred to nitrocellulose membranes, blocked with 10% skim milk in 1× TBST buffer, and then incubated with primary antibodies overnight at 4°C. The following primary antibodies were used: anti-MyHC (Sigma-Aldrich, 05-716-I-100UL, 1:2000), anti-DPF3a (custom made, 1:2000), and anti-GAPDH (Abclonal, AC002, 1:2000). After washing, membranes were incubated with HRP-conjugated secondary anti-mouse IgG (GE, NA931, 1:5000) and anti-rabbit IgG (GE, NA934, 1:5000) for 1 h at room temperature. Signals were visualized using a chemiluminescent HRP substrate (Millipore, WBKLS0500) on a ChemiDoc XRS+ system (Bio-Rad).

### Overexpression in primary myoblasts

Overexpression of DPF3a variants in primary myoblasts was achieved using adenovirus-mediated gene delivery^39^. The cDNA encoding full-length or mutant of DPF3a was cloned into the SalI / EcoRV site of the adenoviral vector pAdTrack-CMV (Addgene, 16405). The resulting vectors were linearized with PmeI (New England Biolabs, R0560S) and then transformed into BJ5183-AD-1 competent cells. Recombinant adenoviral plasmids were transfected into HEK293A cells using Lipofectamine 3000 (Invitrogen, L3000015). The recombinant viruses were propagated in HEK293A cells and purified by CsCl gradient centrifugation. Following infection and removal of the virus, the cells were used for various assays.

### Nucleosome reconstitution

The nucleosome reconstitution was performed with minor modifications to previously described methods ^40,41^. The 65N75-DNA fragment was purified from a PCR product using plasmid pBL842-di-1x-70bp-linker-gal4 (pYM136) as the template and primers #5338 and #primer-Cy5.5 (Supplementary tables S3). The resulting 287-bp product was purified using a Bio-Rad 491 DNA purification system after separation by 3.5% native PAGE. *Xenopus* core histones purified from *E. coli* were used to reconstitute nucleosomes with these DNA products. The H3K36C variant of histone H3 was generated by introducing a lysine-to-cysteine mutation int a vector encoding *Xenopus* H3 C110A, and H3K36me2- and H3K36me3-modified histones were prepared using the methyl-lysine analog (MLA) method^42^. For nucleosome reconstitution, core histones and DNA were mixed at a 1:1.2 molar ratio in TEB buffer (20 mM Tris-HCl pH 7.5, 1 mM EDTA, 1 mM β-mercaptoethanol) containing 2 M NaCl, followed by incubation at 30°C for 2 hours. The mixture was then dialyzed sequentially against decreasing concentrations of NaCl (1.2 M, 1 M, 0.8 M, 0.6 M) at 30°C for 2 hours each, and finally against TEB buffer at 4°C overnight. Mono-nucleosomes were purified using a Bio-Rad 491 system and 3.5% native PAGE, running in 0.3×TBE buffer at 20 W for 2 hours at 4°C. The elution buffer was 10 mM Tris-HCl (pH 7.5). Fractions were analyzed by native PAGE and ethidium bromide staining; nucleosome-containing fractions were pooled and concentrated using a 30-kDa ultrafilter. The concentrated nucleosomes were mixed with storage buffer (250 mM NaCl, 5 mM β-mercaptoethanol, 50% Glycerol) at a 4:1 ratio, and stored at -80°C.

### Recombinant protein purification from *E. coli*

GST-Gal4_DBD_-Flag fusion proteins and GST-HRP2-Flag variants were expressed in BL21 CodonPlus-RIL cells. Cells were cultured in LB medium at 37 °C until the OD_600_ reached 0.4-0.6. Protein expression was induced with 0.2 mM IPTG and the culture was incubated overnight at 16 °C. For lysis, cells were resuspended in 2 M GST buffer (2 M NaCl, 10 mM Tris-HCl pH 7.5, 20 mM Na_3_PO_4_ pH 6.8, 0.01% IGEPAL CA-630, 1 mM β-mercaptoethanol) and lysed by sonication. The lysate was centrifuged at 14,000 rpm for 10 min at 4 °C, and the supernatant was mixed with glutathione agarose beads (Shanghai Chuzhi Bio, SA008010) in 1 M GST buffer (1 M NaCl, 10 mM Tris-HCl pH 7.5, 20 mM Na_3_PO_4_ pH 6.8, 0.01% IGEPAL CA-630, 1 mM β-mercaptoethanol) by adding 0.1 M GST buffer (0.1 M NaCl, 10 mM Tris-HCl pH 7.5, 20 mM Na3PO4 pH 6.8, 0.01% IGEPAL CA-630, 1 mM β-mercaptoethanol). The mixture was incubated for 4 h at 4 °C. After washing, proteins were eluted with GST elution buffer (25 mM Tris-HCl pH 8.0, 50 mM NaCl, 10 mM β-mercaptoethanol, 10% glycerol) containing 100 mM glutathione, and dialyzed against GST elution buffer.

For Gal4_DBD_-Flag fusion proteins and HRP2-Flag variants used in EMSA and remodeling assay, the GST-purified proteins were digested with TEV protease in Flag elution buffer (50 mM HEPES pH 7.9, 100 mM NaCl, 2 mM MgCl_2_, 0.02% IGEPAL CA-630 and 10% glycerol) at 16 °C for 4 h, followed by incubation with anti-DYKDDDDK affinity agarose beads (Smart-Lifesciences, SA042100) overnight at 4 °C. Proteins were then eluted with Flag elution buffer containing 500 μg/mL of 3×FLAG peptide.

### Human cBAF complex purification

Human DPF3a-cBAF complexes were purified from HEK293F cells stably expressing FLAG-tagged DPF3a constructs (as indicated). Cells were washed with cold PBS, and the pellets were resuspended in lysis buffer (10 mM Tris-HCl pH 7.5, 1.5 mM MgCl_2_, 10 mM KCl) freshly supplemented with 1 mM DTT and 1 mM PMSF, followed by rotation for 30 min at 4°C. Nuclei were collected by centrifugation at 4,000 rpm for 5 min at 4°C. The pellets were then resuspended in hypotonic buffer (50 mM Tris-HCl pH 7.5, 1 mM MgCl_2_, 300 mM KCl, 1 mM EDTAand 1% Igepal CA-630) freshly supplemented with a protease inhibitor cocktail (Roche, 04693132001) and rotated for 30 min. The lysates were clarified by centrifugation at 20,000g for 1 h using an SW40Ti rotor (Beckman Coulter). The supernatant was filtered through a 0.45 μm filter and incubated overnight with anti-DYKDDDDK affinity agarose beads (Smart-Lifesciences, SA042100), then eluted with Flag elution buffer (50 mM HEPES pH 7.9, 100 mM NaCl, 2 mM MgCl_2_, 0.02% Igepal CA-630 and 10% glycerol) containing 500 μg/mL of 3×FLAG peptide. The eluted proteins were then subjected to glycerol gradient sedimentation.

### Glycerol gradient sedimentation

Linear 10-30% glycerol gradients were prepared in 14 × 95 mm ultra-clear centrifuge tubes (Beckman Coulter) by layering 10% glycerol solution in Flag elution buffer (50 mM HEPES pH 7.9, 100 mM NaCl, 2 mM MgCl_2_, 0.02% Igepal CA-630 and 10% glycerol) over a 30% glycerol solution (50 mM HEPES pH 7.9, 100 mM NaCl, 2 mM MgCl2, 0.02% Igepal CA-630 and 30% glycerol), followed by horizontal reorientation to establish the density gradient. The FLAG affinity purified DPF3a-cBAF complex (∼ 200 μg) was loaded onto the gradient. The gradients were centrifuged using an SW40Ti rotor at 40,000 rpm for 16 h at 4 °C, and 0.5-mL fractions were collected for silver staining analysis. Fractions containing intact complexes were pooled and concentrated.

### Remodeling assay

Remodeling assays were conducted with minor modifications to previously described methods^43^. Purified cBAF complexes were thawed on ice and diluted in Flag elution buffer. The reaction mixture, also prepared on ice, consisted of remodeling buffer (20 mM Tris-HCl pH 7.5, 50 mM KCl, 5 mM MgCl_2_, 0.1 mg/mL BSA, 10% glycerol), supplemented with designated nucleosomes, 2 mM ATP, and competitor DNA (amounts as indicated). The removal mix contained remodeling buffer, calf thymus DNA and long oligonucleosomes. The reaction was initiated by adding the cBAF complex to the mixture to achieve the specified final concentration, followed by mixing and incubation at 30°C for 1.5 h. After incubation, the removal mix was added, and the mixture was further incubated at 4°C for 1 h. The samples were then loaded onto a 3.5% native polyacrylamide gel (37.5:1) and subjected to electrophoresis in 0.3×TBE buffer at 220 V for 150 minutes at 4°C. The gel was scanned using Li-Cor (LICORbio), and bands were quantified using ImageJ.

### Electrophoretic mobility shift (EMSA) assay

Binding reactions were performed in 15 μL EMSA buffer containing 10 mM HEPES (pH 7.9), 50 mM KCl, 5% glycerol, 4 mM MgCl2, 5 mM DTT, 0.25 mg/mL BSA, and 0.1 mM PMSF. Purified HRP2 protein was incubated with Cy5.5-labeled wild-type, H3K36me2-, or H3K36me3-modified nucleosomes at 30 °C for 45 min. Reaction mixtures were subsequently resolved on 3.5% native polyacrylamide gels (37.5:1 acrylamide:bis-acrylamide) in 0.3× TBE buffer at 220 V for 3 h at 4 °C. Gels were imaged using a Li-COR scanner with the 700 nm detection channel, and band intensities were quantified using ImageJ software.

### GST pull-down assay

Purified GST-HRP2-Flag protein (WT or RR/AA mutant; 0.6 μg) was incubated with pre-equilibrated glutathione Sepharose beads in 100 μL GP300 buffer (50 mM Tris-HCl pH 7.5, 300 mM NaCl, 0.1% Igepal CA-630) at 4 °C for 2 h. Beads were subsequently washed once with GP300 buffer for 5 min at 4 °C and then incubated with 0.45 μg purified cBAF complex at 4 °C for an additional 2 h. Following three washes with GP300 buffer to remove unbound proteins, bead-associated complexes were eluted in 15 μL 1× SDS loading buffer.

### Cleavage Under Targets and Tagmentation (CUT&Tag)

CUT&Tag assays were performed using the Hyperactive Universal CUT&Tag Assay Kit for Illumina (Vazyme, TD903) according to the manufacturer’s instructions. Briefly, 1.5 × 10^5 cells were immobilized on Concanavalin A-coated magnetic beads and incubated overnight at 4 °C with the indicated primary antibody in antibody buffer supplemented with 0.05% digitonin. Cells were subsequently incubated with the corresponding secondary antibody at room temperature for 1 h. After three washes, samples were incubated with hyperactive pA/G-Tn5 transposome at room temperature for 1 h with gentle rotation. Tagmentation was initiated by addition of TTBL buffer and carried out at 37 °C for 1 h. DNA libraries were then amplified and purified using the QIAquick PCR Purification Kit (Qiagen, 28106). Purified libraries were subjected to paired-end 150-bp sequencing on an Illumina NovaSeq platform.

### RNA-seq

Total RNA was isolated from 2 × 10^5^ cells using the RNeasy Mini Kit (Qiagen, 74104) according to the manufacturer’s instructions. Polyadenylated RNA was enriched from 400 ng total RNA using oligo(dT)-conjugated magnetic beads. Strand-specific RNA-seq libraries were generated using the Stranded mRNA-seq Library Prep Kit (Abclonal, RK20349). Library concentrations were quantified using a Qubit fluorometer, and sequencing was performed on an Illumina NovaSeq platform with paired-end 150-bp reads.

### Assay for Transposase-Accessible Chromatin (ATAC-seq)

ATAC-seq was performed as previously described ^44^ with minor modifications. Briefly, 1.5 × 10^5^ primary myoblasts were washed with ice-cold PBS followed by ATAC wash buffer (10 mM Tris-HCl pH 7.4, 10 mM NaCl, 3 mM MgCl2, 0.1% Tween-20). Cells were lysed in ATAC lysis buffer containing 10 mM Tris-HCl pH 7.4, 10 mM NaCl, 3 mM MgCl2, 0.1% Tween-20, 0.01% IGEPAL CA-630, and 0.1% digitonin for 10 min on ice. Nuclei were subsequently washed once with ATAC wash buffer and pelleted by centrifugation. Tagmentation reactions were carried out using the TruePrep DNA Library Prep Kit V2 for Illumina (Vazyme, TD501) at 37 °C for 30 min. Tagmented DNA was amplified using NEBNext High-Fidelity 2× PCR Master Mix (New England Biolabs, M0541S). Libraries were size-selected and purified by agarose gel electrophoresis and subjected to paired-end 150-bp sequencing on an Illumina NovaSeq platform.

### Chromatin Immunoprecipitation sequencing (ChIP-seq)

A total of 1 × 10^6^ cells were dissociated with trypsin and crosslinked in Ham’s F-10 medium containing 1% formaldehyde (Thermo Fisher Scientific, 28908) at room temperature for 10 min. Crosslinking was quenched by addition of glycine to a final concentration of 0.125 M for 10 min at room temperature. Cells were subsequently washed with ice-cold PBS and resuspended in freshly supplemented LB1 buffer (50 mM HEPES-KOH pH 7.5, 140 mM NaCl, 1 mM EDTA, 10% glycerol, 0.5% Igepal CA-630, 0.25% Triton X-100, protease inhibitor cocktail [Roche, 04693132001]). Samples were lysed by vertical rotation at 4 °C for 10 min. Nuclei were pelleted and resuspended in LB2 buffer (10 mM Tris-HCl pH 8.0, 200 mM NaCl, 1 mM EDTA, 0.5 mM EGTA supplemented with protease inhibitors) and incubated with vertical rotation at room temperature for 10 min. Pellets were then resuspended in LB3 buffer (10 mM Tris-HCl pH 8.0, 100 mM NaCl, 1 mM EDTA, 0.5 mM EGTA, 0.1% sodium deoxycholate, 0.5% N-lauroylsarcosine) freshly supplemented with protease inhibitors and 0.1% SDS.

Chromatin was fragmented by sonication using a Branson Sonifier SFX550 for 12 cycles (10 s per cycle, 0.9 s on/0.1 s off). Samples were centrifuged at 15,000 rpm for 10 min at 4 °C, and the supernatant containing sheared chromatin was collected. Triton X-100 and SDS were adjusted to final concentrations of 1% and 0.1%, respectively, prior to overnight incubation with the indicated antibodies at 4 °C with rotation. The following antibodies were used for ChIP assays: H3K36me2 (Abcam, ab9049), total H3 (Abcam, ab1791), and CTCF (Cell Signaling Technology, 3418S).

For immunoprecipitation, 30 μL magnetic Protein G Dynabeads slurry (Invitrogen, 10004D) was pre-washed three times with blocking buffer (PBS containing 5 mg/mL BSA) and subsequently added to the chromatin-antibody mixtures for incubation at 4 °C for 4 h with rotation. Beads were then washed sequentially with low-salt wash buffer (20 mM Tris-HCl pH 8.1, 2 mM EDTA, 150 mM NaCl, 1% Triton X-100, 0.1% SDS), high-salt wash buffer (20 mM Tris-HCl pH 8.1, 2 mM EDTA, 500 mM NaCl, 1% Triton X-100, 0.1% SDS), LiCl wash buffer (10 mM Tris-HCl pH 8.1, 1 mM EDTA, 250 mM LiCl, 1% NP-40, 1% sodium deoxycholate), and twice with TE buffer (10 mM Tris-HCl pH 8.1, 1 mM EDTA, 50 mM NaCl).

DNA-protein complexes were eluted in 200 μL elution buffer (50 mM Tris-HCl pH 8.0, 10 mM EDTA, 1% SDS) at 65 °C for 15 min. Crosslinks were reversed by addition of 8 μL 5 M NaCl followed by incubation at 65 °C for 4 h. Samples were then treated with RNase A (Takara, 2158) at 37 °C for 30 min and subsequently incubated with 4 μL 0.5 M EDTA, 8 μL 1 M Tris-HCl (pH 6.5), and 1 μL 10 mg/mL Proteinase K (New England Biolabs, P8107S) at 45 °C for 1.5 h. DNA was purified using the QIAquick PCR Purification Kit (Qiagen, 28106). Sequencing libraries were prepared using the VAHTS Universal DNA Library Prep Kit for Illumina V3 (Vazyme, ND607) and subjected to paired-end 150-bp sequencing on an Illumina NovaSeq platform.

### RNA-seq analysis

Raw sequencing reads were subjected to adapter trimming and quality filtering using fastp/0.23.2 with the following parameters: -g, -q 5, -u 50, -n 15, -l 150, -overlap_diff_limit 1, and -- overlap_diff_percent_limit 10. Quality control metrics were assessed via fastp-generated reports to ensure effective removal of low-quality sequences. High-quality reads were aligned to the mouse reference genome (mm39/GRCm39) using STAR (v.2.7.10a)^45^ with the parameters --outSAMtype BAM SortedByCoordinate --twopassMode Basic --outFilterMismatchNmax 2 --outSJfilterReads Unique --quantMode TranscriptomeSAM GeneCounts. Duplicates were marked and removed with Picard (v.3.1.0). Next, the featureCounts function of the subread (v.2.0.1)^46^ was used to assign reads to genes. Assigned reads were then normalized, and differential expression analysis was performed using the R package edgeR (v.4.4.2)^47^. Differentially expressed genes (DEGs) were called using an adjusted *P* value ≤ 0.05 and |log_2_FC| ≥ 1. Volcano plots were generated with the R package ggplot2 (v.3.5.1) to visualize edgeR results. FPKM values for gene expression were calculated using RSEM (v.1.3.3)^48^. RPKM-normalized coverage profiles were generated with deepTools (v.3.5.1)^49^. Gene Ontology (GO) enrichment analysis was performed on significantly up- or down-regulated genes using the R package clusterProfiler (v.4.12.6). GO Biological Processes categories were selected, and adjusted *P* values for the top terms were visualized as bubble plots.

### ATAC-seq analysis

Raw ATAC-seq reads were trimmed with fastp/0.23.2 and aligned to the mouse reference genome (mm39/GRCm39) using Bowtie2 (v.2.2.5)^50^ with the parameters --end-to-end --very-sensitive --no-mixed --no-discordant -X 1000. PCR duplicates were removed using Picard (v.3.1.0). Mitochondrial contamination was removed using samtools^51^. Peaks were called using MACS3 (v.3.0.1)^52^. bigWig files were generated for visualization from individual and merged BAM files using Deeptools^49^. Heatmaps and average profiles were generated from merged bigWig files using Deeptools.

Differentially accessible peaks were identified by comparing *Dpf3a*KO samples to wild-type controls using EdgeR (v.4.4.2). ATAC-seq peaks with an FDR < 0.05 were considered significantly gained or reduced. Metagene profiles and heatmaps showing changes in chromatin accessibility were generated using deepTools (v.3.5.1)^49^. For motif enrichment analysis of ATAC-seq clusters, known motifs were identified using Homer (v.4.11) findMotifsGenome.pl with the parameter -size 200.

### CUT&Tag and ChIP-seq analysis

Raw sequencing reads were trimmed with fastp/0.23.2 and aligned to the mouse reference genome (mm39/GRCm39) using Bowtie2 (v.2.2.5)^50^ with the parameters --end-to-end --very-sensitive --no-mixed --no-discordant -X 1000. PCR duplicates were removed using Picard (v.3.1.0). bigWig files were generated for visualization from individual and merged BAM files using Deeptools^49^ bamCoverage with the parameters --normalizeUsing RPKM --binsize 1. For DPF3a visualization, the signals were obtained after subtracting non-specific signals present in the *Dpf3a* KO cells. EdgeR (v.4.4.2) was employed to compare ATAC-seq signals at DPF3a-bound regions between WT and KO samples. ATAC-seq peaks were considered gained or reduced as calculated by edgeR (FDR < 0.05). Heatmaps and average profiles were generated from merged bigWig files using Deeptools computeMatrix followed by plotHeatmap, with the center of the peaks as the reference point. To precisely quantify ATAC-seq, H3K36me3 and H3 read intensity at DPF3a regions, the matrix containing scores per genomic region calculated with computeMatrix was exported and plotted in R. The ChIPSeeker package (v.1.40.0) ^53^ was used to annotate peaks and determine their feature distribution with the annotation package TxDb.Mmusculus.UCSC.mm39.knownGene. Chromatin state modeling was performed using the ChromHMM software ^54,55^ as previously described. The overlap between each state and DPF3a peaks was calculated using bedtools annotate.

### Chromatin Conformation Capture (3C) assay

The chromosome conformation capture (3C) assay was performed based on a previously described protocol ^56^ with minor modifications. Briefly, 1 × 10^6^primary myoblasts were dissociated with trypsin and crosslinked in 2% formaldehyde (Thermo Fisher Scientific, 28908) with rotation at room temperature for 10 min. Crosslinking was quenched by addition of glycine to a final concentration of 125 mM followed by rotation at room temperature for an additional 10 min. Cells were washed twice with ice-cold PBS and lysed in freshly prepared lysis buffer containing 10 mM Tris-HCl pH 7.4, 0.5% Igepal CA-630, 1% Triton X-100, 150 mM NaCl, and 5 mM EDTA supplemented with protease inhibitor cocktail (Roche, 04693132001).

Isolated nuclei were washed once with 450 μL 1.2× NEBuffer DpnII (New England Biolabs, R0543S) and resuspended in 500 μL 1.2× NEBuffer DpnII supplemented with 15 μL 10% SDS (final concentration 0.3%). Samples were incubated at 37 °C with shaking for 1 h to permeabilize chromatin. Triton X-100 was subsequently added to a final concentration of 2.5% to sequester SDS, followed by incubation at 37 °C with shaking for an additional 1 h. Chromatin was digested with 100 U DpnII (New England Biolabs, R0543S) at 37 °C for 1 h with shaking, followed by a second digestion with an additional 100 U DpnII overnight. Digestion efficiency was verified by agarose gel electrophoresis on a 0.6% gel. DpnII was heat-inactivated at 65 °C for 20 min.

Digested chromatin was transferred to 15 mL tubes and ligated in a reaction containing 700 μL 10× T4 DNA Ligase Reaction Buffer (New England Biolabs, B0202S) and 1200 U T4 DNA ligase (New England Biolabs, M0202M). The total reaction volume was adjusted to 7 mL with nuclease-free H2O, and ligation was performed overnight at 16 °C. Protein digestion was subsequently carried out by addition of 30 μL Proteinase K (New England Biolabs, P8107S) followed by incubation overnight at 16 °C. Ligated DNA products were purified using the QIAquick PCR Purification Kit (Qiagen, 28106) and quantified by qPCR using the indicated primer sets (Supplementary Table S3).

### Statistical analysis

Statistical analyses were performed using the GraphPad Prism software (v.11.0.1) and statistical details are provided in figure legends. Significance was defined by *P* values determined by a two-tailed unpaired Student’s *t*-test.

## Results

### DPF3a orchestrates skeletal muscle regeneration through the regulation of myogenic differentiation

The mammalian canonical BAF (cBAF; BRG1/BRM-associated factor) complex contains multiple non-catalytic regulatory subunits that modulate chromatin remodeling activity and confer functional specificity. Among these, the DPF (double PHD finger) family proteins: DPF1, DPF2, and the two DPF3 isoforms, DPF3a and DPF3b, are incorporated as mutually exclusive monomeric subunits within the complex^15^. DPF3a is structurally distinguished from other DPF family members by a truncated PHD domain and a unique C-terminal region, suggesting specialized functions beyond those of the core remodeling machinery. Consistent with this notion, Dpf3 transcripts exhibit pronounced tissue specificity, with enriched expression in skeletal muscle, heart, testis, and ovary (Supplementary Fig. S1a). Previous studies identified DPF3a as a cBAF-associated factor capable of interacting with HRP2 to promote transcriptional activation^27^, and genetic studies in zebrafish implicated DPF3 proteins in muscle development^24^. However, the physiological function of DPF3a in mammalian tissues, as well as its mechanistic contribution to chromatin regulation, has remained incompletely understood.

To define the specific role of DPF3a independently of DPF3b, we generated a Dpf3a-selective knockout mouse model using CRISPR-Cas9-mediated genome editing. DPF3a and DPF3b share exons 1-8, which encode the N-terminal 291 amino acids, whereas the DPF3a-specific C-terminal region is encoded by exon 9. We therefore targeted exon 9 to selectively disrupt DPF3a while preserving DPF3b expression, generating a truncated DPF3 protein (DPF3t) lacking the DPF3a-specific C-terminal domain (Fig. 1a). Genotyping PCR confirmed successful targeting, and immunoblotting using an antibody directed against the DPF3a C-terminus verified complete loss of DPF3a protein in primary myoblasts derived from knockout mice (Fig. 1b and Supplementary Fig. S1b). Dpf3a KO mice were viable, fertile, and grossly indistinguishable from wild-type littermates, indicating that DPF3a is dispensable for embryonic development and organismal viability under homeostatic conditions.

**Figure 1.**
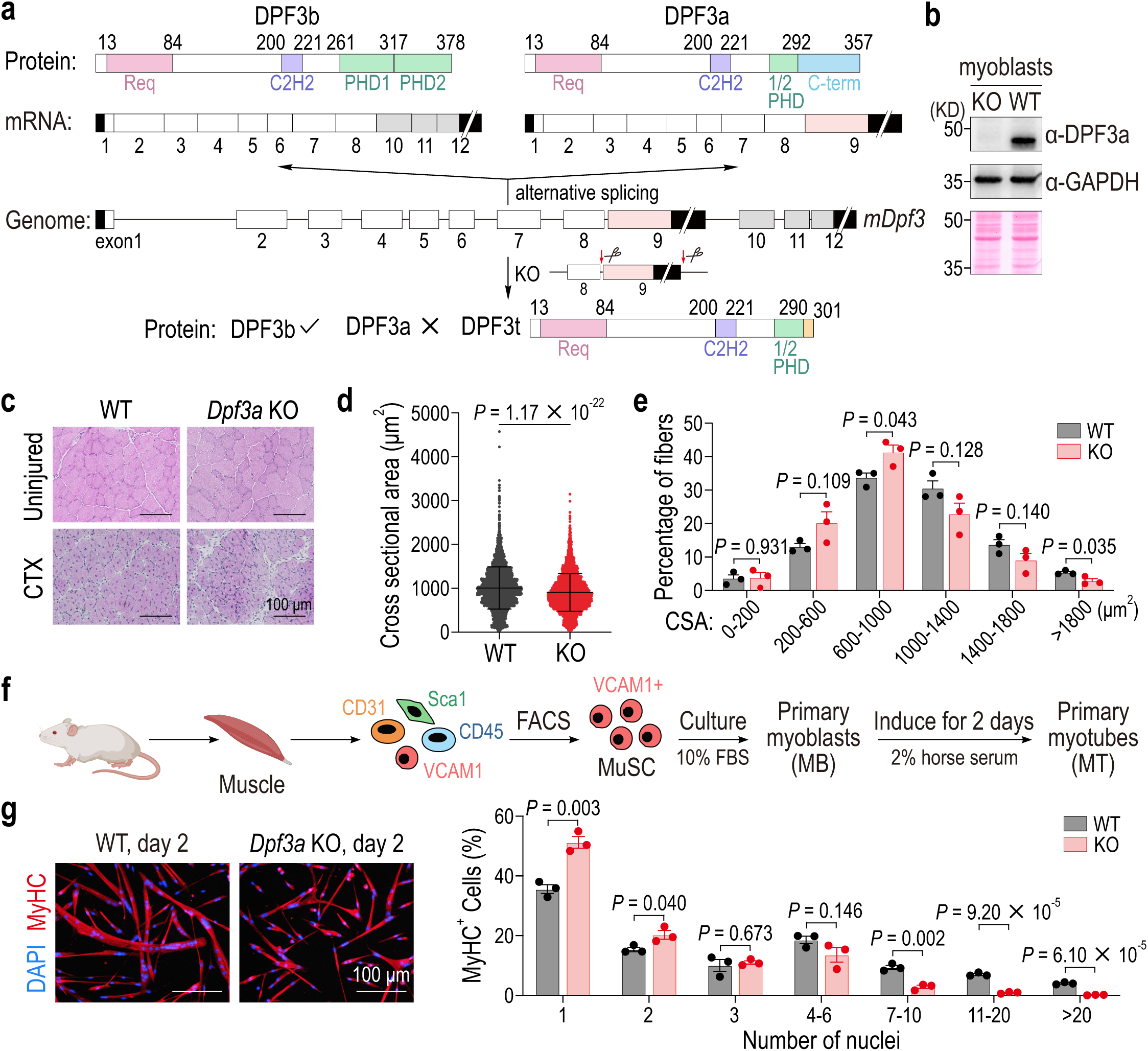
DPF3a orchestrates skeletal muscle regeneration through the regulation of myogenic differentiation. **a.** Strategy for generating *Dpf3a* knockout mice. From top to bottom: protein structures of DPF3b and DPF3a isoforms; mRNA of *Dpf3b* and *Dpf3a* with exon organization; genomic locus of the *Dpf3* gene encoding both isoforms, with the locations of designed sgRNAs (indicated by scissors). DPF3t, truncated DPF3 protein. **b.** Validation of *Dpf3a*knockout mice in myoblasts derived from muscle tissue by western blot using whole-cell lysate. The DPF3a antibody specifically recognizes the C-terminal domain of DPF3a. **c.** Representative view of hematoxylin and eosin staining of tibialis anterior (TA) muscle cross sections from wild-type (WT) and *Dpf3a* knockout (KO) mice at 7 days after injury. **d.** Quantification of myofiber cross-sectional area (CSA) to assess muscle regeneration; data are shown as mean ± s.d. of all measured myofibers from three WT (n = 3699) and three *Dpf3a* KO mice (n = 3852). **e.** Distribution of myofiber cross-sectional area (CSA) in WT and *Dpf3a* KO mice. **f.** Schematic of the *in vitro* differentiation from primary myoblasts to myotubes. **g.** (Left) Representative view of immunofluorescence staining of myosin heavy chain (MyHC) and nuclei (DAPI) in WT and *Dpf3a*KO cells after 2 days of differentiation. (Right) Quantification of multinucleation in MyHC-positive cells from three independent experiments; data are shown as mean ± s.e.m.

We next examined whether DPF3a contributes to skeletal muscle regeneration *in vivo*. Tibialis anterior (TA) muscles of wild-type (WT) and Dpf3a KO mice were subjected to cardiotoxin (CTX)-induced injury, and regenerative responses were analyzed seven days post-injury. Histological examination revealed centrally nucleated regenerating myofibers in both genotypes, indicating successful initiation of the regenerative program (Fig. 1c). However, regenerating myofibers in Dpf3a KO muscle exhibited a marked reduction in cross-sectional area (CSA) relative to WT controls (Fig. 1c-d). Consistently, fiber size distribution analysis demonstrated a substantial depletion of large regenerating myofibers (>1800 μm^2^) accompanied by a corresponding enrichment of smaller fibers in Dpf3a KO muscle (Fig. 1e). These findings establish DPF3a as an important regulator of efficient skeletal muscle regeneration.

To determine whether the regenerative phenotype reflects a cell-autonomous function in muscle progenitors, we isolated primary muscle stem cells (MuSCs) from WT and Dpf3a KO mice by fluorescence-activated cell sorting (FACS) using established surface markers (VCAM1+Sca1−CD31−CD45−; Supplementary Fig. S1c-d). Purified MuSCs were expanded as primary myoblasts and induced to differentiate by serum withdrawal (Fig. 1f). Following two days of differentiation, WT myoblasts robustly fused into elongated multinucleated myotubes, whereas Dpf3a-deficient cells formed markedly smaller and shorter myotubes with impaired multinucleation (Fig. 1g). Quantitative analysis further revealed a significant increase in the proportion of MyHC-positive mononucleated cells and a concomitant reduction in highly multinucleated myotubes containing seven or more nuclei in Dpf3a KO cultures (Fig. 1g). These observations demonstrate that DPF3a functions intrinsically within myogenic progenitors to promote myoblast fusion and terminal differentiation.

Interestingly, comparison with previously reported Hrp2-deficient mice revealed partially overlapping yet distinct regenerative phenotypes^27^. Whereas HRP2 loss predominantly affected smaller regenerating fibers, Dpf3a deficiency preferentially reduced the abundance of larger myofibers (Fig. 1e). This distinction may reflect the broader expression pattern and pleiotropic functions of HRP2 relative to the more tissue-restricted expression of DPF3a. Nonetheless, the phenotypic convergence supports the possibility that DPF3a and HRP2 operate within a shared regulatory pathway during myogenesis, with DPF3a acting as a specialized cBAF-associated effector, a model explored in subsequent mechanistic analyses.

### DPF3a acts as a central transcriptional co-regulator of the myogenic gene expression program

Myogenesis is driven by a highly coordinated transcriptional program that integrates lineage-specific transcription factors with dynamic chromatin regulation^30^. The pronounced differentiation defects observed in Dpf3a-deficient myoblasts prompted us to investigate the transcriptional programs governed by DPF3a during myogenic progression. We hypothesized that DPF3a promotes muscle differentiation through broad regulation of myogenic gene expression downstream of the cBAF chromatin remodeling machinery. To examine this possibility, we performed high-resolution temporal RNA-seq profiling of WT and Dpf3a KO primary myoblasts across a three-day differentiation time course (Fig. 2a).

**Figure 2.**
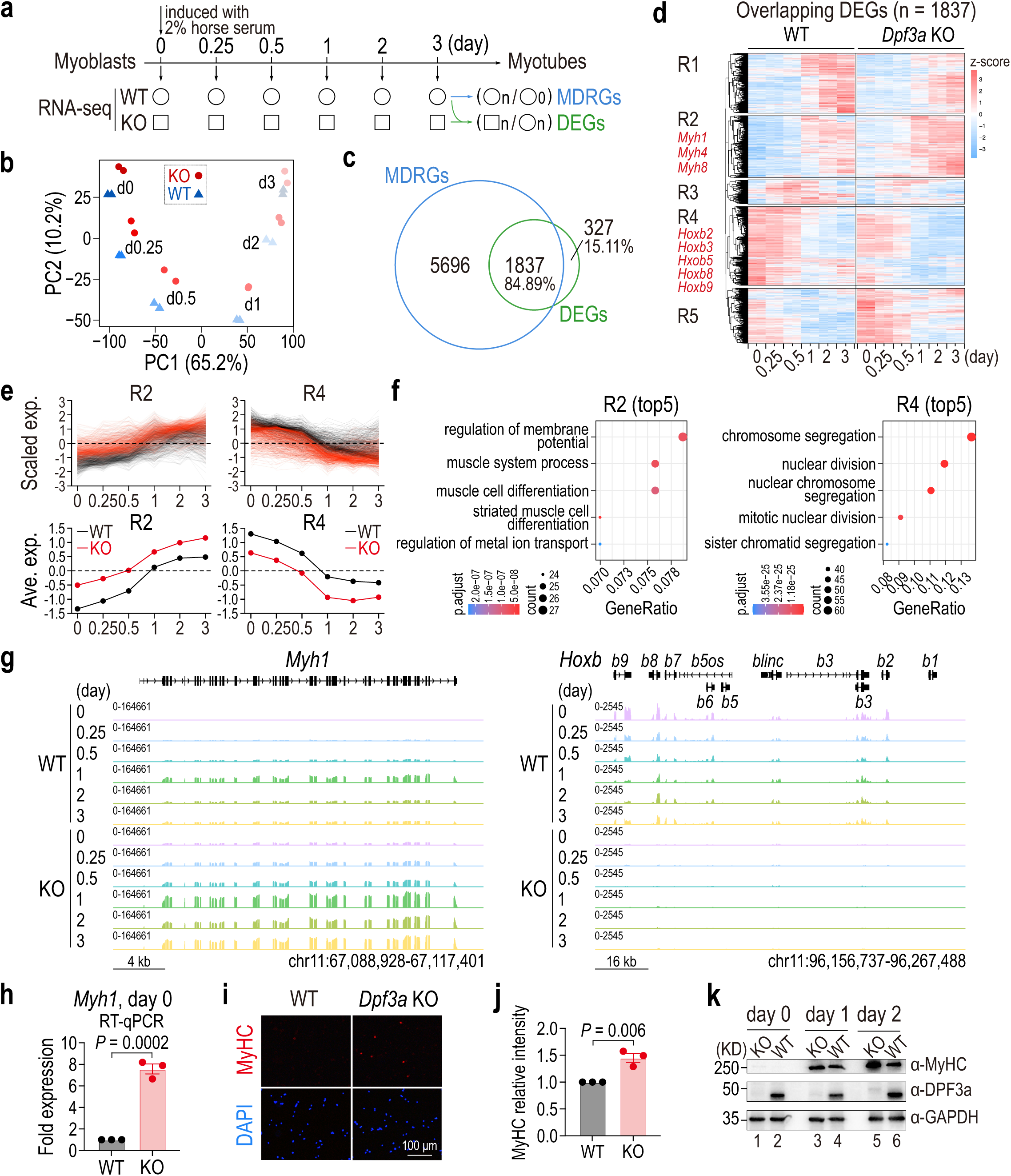
DPF3a acts as a central transcriptional co-regulator of the myogenic gene expression program. **a.** Schematic of the experimental strategy to identify muscle differentiation-related genes (MDRGs) and DPF3a-dependent differentially expressed genes (DEGs) by multi-timepoint RNA-seq of differentiating WT and *Dpf3a* KO myoblasts. **b.** Principal component analysis (PCA) of the multi-timepoint RNA-seq profiling during myoblast differentiation from two biological replicates. **c.** Venn diagram showing the overlap between MDRGs and DPF3a-dependent DEGs. **d.** Heatmap illustrating the expression patterns of the overlapping DEGs across all time points in WT and *Dpf3a* KO cells. Each column represents a biological replicate. **e.** (Top) Scaled expression (Scaled exp.) and (bottom) average expression (Ave. exp.) dynamics of the R2 and R4 cluster genes identified in **d**. **f.** Gene Ontology (GO) analysis (Biological Processes) of the R2 and R4 cluster genes shown in **d**. **g.** Genome browser tracks of RNA-seq signals at the *Myh1* and *Hoxb* loci in WT and *Dpf3a* KO myoblasts. **h.** Expression levels of *Myh1* in WT and *Dpf3a* KO myoblasts as assessed by RT-qPCR from three independent experiments; data are shown as mean ± s.e.m. **i.** Representative view of immunofluorescence staining of MyHC (encoded by *Myh1*) in WT and *Dpf3a* KO myoblasts. **j.** Quantification of the relative intensity of MyHC in WT and *Dpf3a* KO myoblasts from three independent experiments. **k.** Western blot analysis of MyHC protein expression dynamics during myoblast differentiation (days 0, 1 and 2) in WT and *Dpf3a* KO cells.

Principal component analysis (PCA) revealed a continuous developmental trajectory in WT samples that closely mirrored morphological differentiation, whereas Dpf3a KO samples diverged progressively from the WT trajectory at each corresponding time point (Fig. 2b). These findings indicate that loss of DPF3a results in pervasive and sustained transcriptional perturbation throughout myogenic differentiation. Notably, expression of Myod1, the master myogenic determinant responsible for initiating the differentiation program^57,58^, remained comparable between WT and KO cells (Supplementary Fig. S2a-b). Thus, DPF3a is not required to establish myogenic competency per se, but instead appears to fine-tune the fidelity, timing, and stability of downstream transcriptional programs during differentiation.

To systematically define the DPF3a-dependent transcriptional network, we first identified muscle differentiation-related genes (MDRGs) based on genes exhibiting significant temporal expression changes during WT differentiation. In parallel, we defined DPF3a-dependent differentially expressed genes (DEGs) by aggregating genes displaying significant genotype-dependent differences at any differentiation stage. Strikingly, the majority of DEGs overlapped with the MDRG set (Fig. 2c), indicating that DPF3a predominantly regulates genes intrinsically associated with the myogenic differentiation program rather than unrelated transcriptional pathways. Hierarchical clustering of the 1,837 overlapping genes resolved five major expression modules (R1-R5) defined by both temporal dynamics and genotype-specific regulation (Fig. 2d-e and Supplementary Fig. S2c). R1 comprised progressively induced genes with higher expression in WT cells, whereas R2 contained genes that similarly increased during differentiation but were aberrantly elevated in Dpf3a KO cells. R3 represented transiently induced genes that subsequently declined, while R4 and R5 consisted of progressively downregulated genes preferentially enriched in WT and KO cells, respectively. Importantly, several canonical fast myosin heavy chain genes, including Myh1, Myh4, and Myh8, clustered within R2, whereas multiple members of the Hoxb cluster, including Hoxb2, Hoxb3, Hoxb5, Hoxb8, and Hoxb9, were assigned to R4. These expression patterns closely matched previous reports^59^, supporting the robustness of the clustering analysis. Gene ontology analysis further revealed that R2 was strongly enriched for muscle system processes, whereas R4 was associated with mitotic cell cycle regulation (Fig. 2f), suggesting that DPF3a coordinates both activation of differentiation-associated genes and repression of proliferative programs. Additional clusters encompassed genes linked to diverse biological processes, indicating a broader role for DPF3a in maintaining transcriptional homeostasis during myogenesis (Supplementary Fig. S2d).

Examination of individual genomic loci further revealed that DPF3a regulates transcription in a highly coordinated regional manner. The fast myosin heavy chain (fMyh) locus, which encodes major contractile components of mature skeletal muscle^60^, exhibited precocious and exaggerated activation in Dpf3a KO cells, whereas the developmentally regulated Hoxb cluster showed widespread transcriptional repression (Fig. 2g and Supplementary Fig. S2e). Notably, genes within each locus displayed concordant directional changes, while distinct loci responded oppositely, suggesting that DPF3a regulates transcription across defined chromosomal domains rather than at isolated single-gene targets or through broad genome-wide effects. To further examine the role of DPF3a in maintaining transcriptional restraint prior to differentiation, we analyzed Myh1 expression in proliferating myoblasts. RT-qPCR demonstrated substantial upregulation of Myh1 transcripts in Dpf3a KO cells relative to WT controls (Fig. 2h). Consistently, both immunofluorescence staining and immunoblotting revealed markedly elevated MyHC protein levels in Dpf3a-deficient myoblasts under steady-state conditions (Fig. 2i-j) and throughout early differentiation (days 0-2; Fig. 2k). These findings indicate that DPF3a is required to suppress premature activation of fast myosin genes in undifferentiated myoblasts, thereby preserving appropriate temporal progression of the differentiation program.

Interestingly, DPF3a itself displayed dynamic temporal regulation during myogenesis. In WT cells, Dpf3a transcripts peaked at day 1 of differentiation and subsequently declined, whereas DPF3a protein accumulation peaked at day 2, consistent with delayed protein accumulation following transcriptional induction (Supplementary Fig. S2f-h). This temporally phased expression pattern suggests that DPF3a activity is tightly synchronized with specific stages of myogenic progression. Importantly, Dpf3b expression remained unchanged in Dpf3a KO cells (Supplementary Fig. S2i), confirming that the observed phenotypes arise specifically from DPF3a loss rather than compensatory dysregulation of the alternative isoform.

Collectively, these results establish DPF3a as a temporally regulated transcriptional co-regulator that maintains the fidelity of myogenic gene expression programs, in part by preventing premature activation of differentiation-associated loci and coordinating transcriptional regulation across discrete chromosomal domains.

### DPF3a preferentially occupies focal depressions embedded within broad H3K36me2 chromatin domains

Acentral unresolved question concerns how DPF3a achieves selective genomic localization. Because transcriptional perturbations were already evident in undifferentiated Dpf3a KO myoblasts (Supplementary Fig. S2j-k), we focused subsequent mechanistic analyses on the myoblast state to define the chromatin basis of DPF3a function prior to overt differentiation defects. To map DPF3a occupancy genome-wide, we performed CUT&Tag profiling using WT and Dpf3a KO primary myoblasts, with knockout samples serving as controls to remove non-specific signals. Genomic annotation revealed that DPF3a binding sites were predominantly located within distal intergenic regions (29.39%), intronic regions (37.38%), and promoters (≤1 kb; 23.02%) (Fig. 3a), consistent with the established role of cBAF complexes at enhancer-associated regulatory elements^61^. To investigate whether DPF3a binding correlates with local remodeling activity, we additionally performed ATAC-seq in WT and Dpf3a KO myoblasts. Based on accessibility changes at DPF3a-bound regions, sites were classified into three groups: G1 regions exhibiting increased accessibility in KO cells (n=2,011), G2 regions exhibiting reduced accessibility (n=1,988), and G3 regions displaying minimal accessibility changes (Fig. 3b). These groups provided a framework for subsequent epigenomic characterization.

**Figure 3.**
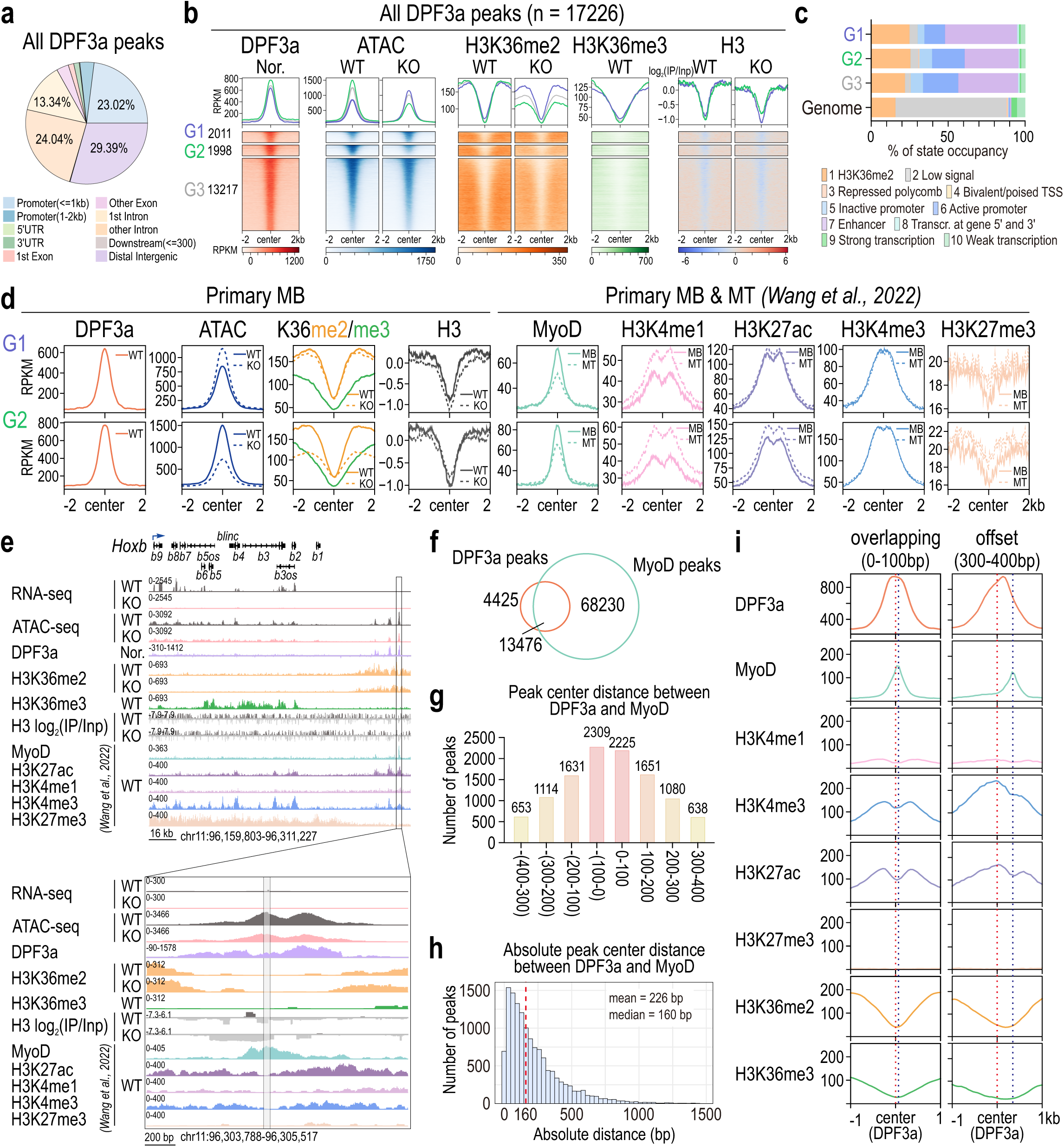
DPF3a preferentially occupies focal depressions embedded within broad H3K36me2 chromatin domains. **a.** Pie chart depicting the genomic annotation distribution of all DPF3a binding peaks. **b.** Heatmap illustrating H3K36me2/3 modification patterns at DPF3a binding regions, categorized into G1, G2 and G3 clusters based on distinct accessibility dynamics. ‘Nor.’ indicates normalized DPF3a signal generated by subtracting non-specific signal present in the *Dpf3a* KO myoblasts. **c.** Distribution of chromatin states of the G1, G2, and G3 cluster regions. **d.** Meta-plot showing the distribution of epigenetic signals in G1 and G2 regions. MB, myoblast; MT, myotube. **e.** Representative genome browser tracks at the *Hoxb* genomic locus in WT and *Dpf3a* KO myoblasts. Top, the entire *Hoxb* locus; bottom, zoom-in view of the region indicated by the box on the top. **f.** Venn diagrams showing the overlap between DPF3a and MyoD binding peaks. **g.** Distribution of peak center distance between DPF3a and MyoD binding peaks. **h.** Distribution of absolute peak center distance between DPF3a and MyoD binding peaks. The red line indicates the median value. **i.** Meta-plot showing the distribution of epigenetic signals at DPF3a binding peaks (from −1 kb to the center to +1 kb), focusing on the overlapping (0-100 bp) and offset (300-400 bp) intervals relative to the peak center distance, as defined in **h**.

To define the chromatin context surrounding DPF3a occupancy, we profiled H3K36me2 and H3K36me3 by ChIP-seq. Correlation analyses demonstrated high reproducibility across biological replicates for both histone modification and accessibility datasets (Supplementary Fig. S3a). Genome-wide H3K36me2/3 enrichment patterns across gene bodies were consistent with previously established chromatin landscapes (Supplementary Fig. S3b), and comparison with published mouse embryonic stem cell datasets^29^ further validated the quality and specificity of our data, including at lineage-relevant loci such as Myod1 (Supplementary Fig. S3c).

Unexpectedly, DPF3a occupancy did not simply coincide with maximal H3K36me2 enrichment. Instead, DPF3a preferentially localized to focal regions of relative H3K36me2 depletion embedded within broad H3K36me2-enriched domains, a distinctive chromatin configuration that we term depressions within H3K36me2 plateaus (Fig. 3b). Heatmap analyses stratified by accessibility dynamics confirmed that this architecture was consistently associated with DPF3a-bound regions across all accessibility classes (Fig. 3b). ChromHMM integration with published histone modification datasets, including H3K27ac, H3K4me1, H3K4me3, and H3K27me3, further demonstrated that DPF3a-associated regions are strongly enriched for H3K36me2 signatures relative to the genomic background while simultaneously exhibiting active promoter and enhancer chromatin states (Fig. 3c and Supplementary Fig. S3d). Meta-profile analyses revealed an inverse relationship between DPF3a occupancy and local H3K36me2 intensity, accompanied by sharp enrichment of MyoD and H3K4me1 signals at DPF3a-centered regions (Fig. 3d and Supplementary Fig. S3e). Notably, MyoD occupancy was more prominent in myoblasts, whereas H3K4me1 enrichment increased during myotube differentiation, suggesting that these regions represent dynamically regulated regulatory elements undergoing chromatin state transitions during myogenesis.

Genome browser inspection of representative loci further illustrated this unique chromatin architecture. At both the fast myosin heavy chain (fMyh) and Hoxb loci, DPF3a peaks localized precisely within focal depressions nested inside broad H3K36me2 domains (Fig. 3e and Supplementary Fig. S3f). These regions frequently coincided with nucleosome-depleted intervals marked by accessible ATAC-seq signals and were enriched for active chromatin marks including H3K27ac, H3K4me1, and H3K4me3, while displaying local depletion of H3K27me3. Collectively, these features define a poised yet active regulatory chromatin environment associated with DPF3a occupancy.

The widespread genomic distribution of H3K36me2 raised the question of how DPF3a-cBAF achieves positional specificity within such extensive chromatin territories. The coincident enrichment of MyoD and H3K4me1 at DPF3a-bound regions suggested that lineage-specific transcription factors may provide an additional layer of targeting specificity. Consistent with this model, overlap analysis revealed that 13,476 DPF3a peaks coincided with MyoD occupancy, whereas only 4,425 peaks were uniquely associated with DPF3a (Fig. 3f). These findings indicate that DPF3a targeting is strongly coupled to the core myogenic transcriptional circuitry.

Closer examination of the spatial relationship between DPF3a and MyoD uncovered a highly ordered chromatin arrangement. Analysis of peak-center distances demonstrated that the majority of overlapping DPF3a-MyoD peaks were separated by less than 100 bp, with progressively fewer peaks distributed across increasing distance intervals (Fig. 3g). The overall mean center-to-center distance was approximately 226 bp (Fig. 3h), closely corresponding to the span of a nucleosome together with linker DNA. Comparison of closely overlapping versus spatially offset peaks revealed corresponding positional shifts in both MyoD and histone modification profiles relative to DPF3a occupancy (Fig. 3i). Interestingly, although these regions frequently displayed reduced accessibility in Dpf3a KO cells (Supplementary Fig. S3g), associated genes exhibited only limited transcriptional perturbation (Supplementary Fig. S3h-i), implying that DPF3a-dependent regulation at these sites may not primarily operate through direct modulation of promoter accessibility.

Together, these findings define a distinctive targeting principle for DPF3a-cBAF. DPF3a preferentially occupies focal depressions embedded within broad H3K36me2 plateaus, where relative H3K36me2 depletion coincides with lineage-specific transcription factor binding and active regulatory chromatin features. This organization suggests a hierarchical targeting mechanism in which the H3K36me2 landscape establishes broad permissive chromatin territories, while MyoD and associated regulatory factors refine precise recruitment sites. The characteristic ∼226 bp spatial offset between DPF3a and MyoD further raises the possibility that DPF3a-cBAF engages nucleosomes adjacent to transcription factor binding sites in a spatially constrained manner, thereby enabling selective remodeling within complex regulatory chromatin environments.

### Association with the H3K36me2/3 reader HRP2 is essential for DPF3a function *in vivo*

The distinctive localization pattern of DPF3a within H3K36me2-enriched chromatin domains suggested that additional molecular interactions are required to confer genomic specificity. Previous studies demonstrated that DPF3a interacts with HRP2, a PWWP-domain protein capable of recognizing H3K36me2 and H3K36me3 modifications^26,27^. However, whether this interaction is functionally required for DPF3a activity in primary myoblasts remained unresolved. To directly test the importance of HRP2 association, we generated a DPF3a mutant harboring F325A/F328A substitutions (FF/AA) within the C-terminal region, mutations previously shown to disrupt HRP2 binding^27^ (Fig. 4a). We then compared the ability of full-length (FL) and FF/AA mutant DPF3a to rescue the differentiation defects of Dpf3a KO primary myoblasts.

**Figure 4.**
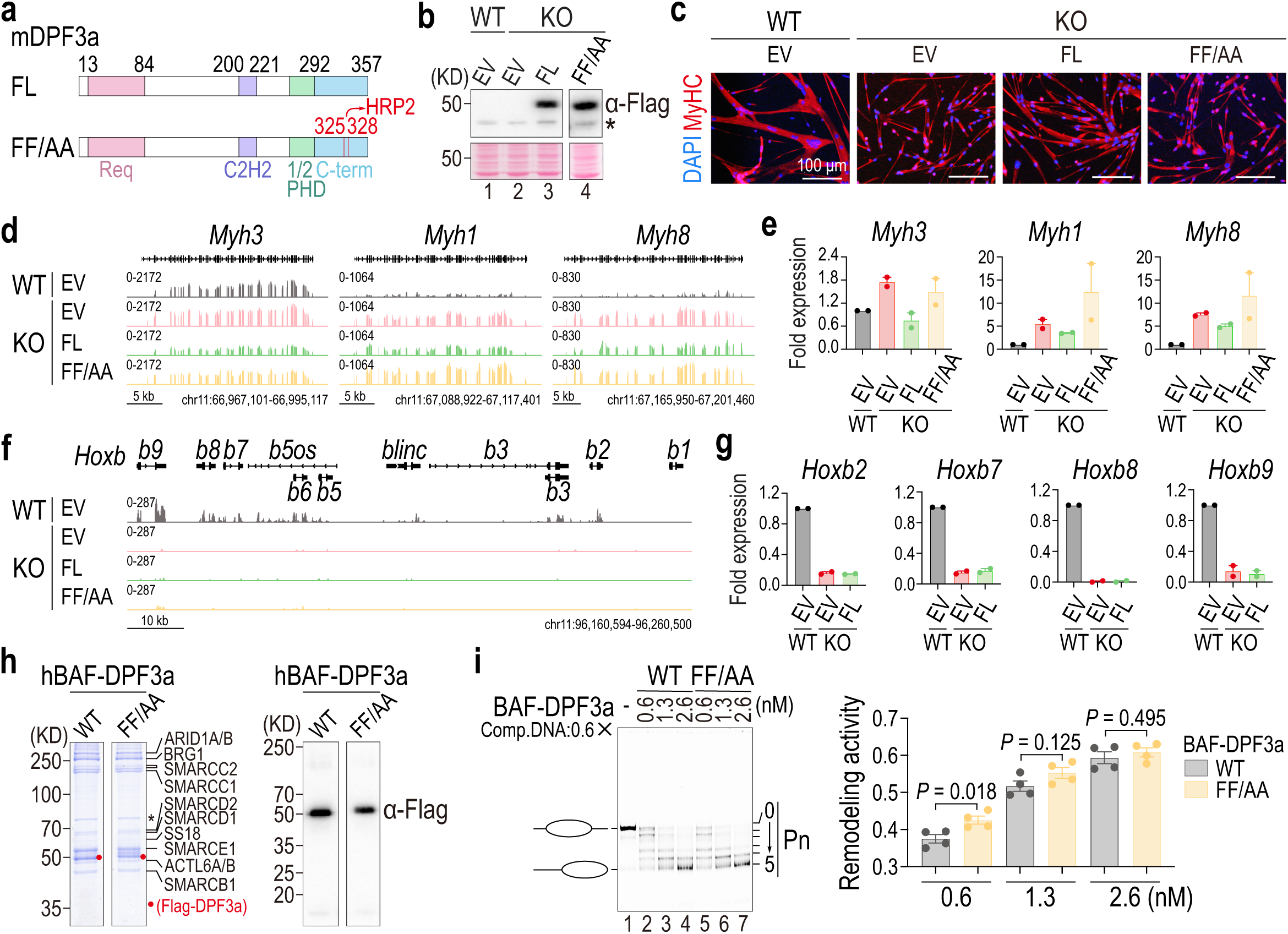
Association with the H3K36me2/3 reader HRP2 is essential for DPF3a function in vivo. **a.** Schematic of full-length (FL) and FF/AA mutant (HRP2-binding defective) mouse DPF3a proteins; FF/AA, F325A/F328A double mutation in DPF3a that disrupts interaction with HRP2, showed as red lines. **b.** Detection of exogenously expressed FL and FF/AA DPF3a in *Dpf3a* KO myoblasts by western blot using whole cell lysate; asterisks indicate non-specific bands. **c.** Representative view of immunofluorescence staining of myosin heavy chain (MyHC) and nuclei (DAPI) in cells overexpressing empty vector (EV), FL or FF/AA DPF3a after 2 days differentiation. **d.** Representative genome browser tracks of RNA-seq at *Myh3*, *Myh1* and *Myh8*in WT (EV) and KO (EV, FL and FF/AA) myoblasts. **e.** RT-qPCR analysis of *Myh3*, *Myh1* and *Myh8* in the same samples as in **d**. **f.** Representative genome browser tracks of RNA-seq at the *Hoxb* locus in WT (EV) and KO (EV, FL and FF/AA) myoblasts. **g.** RT-qPCR analysis of *Hoxb2*, *Hoxb7*, *Hoxb8* and *Hoxb9* in the same samples as in **f**. **h.** Coomassie blue staining (left) and western blot (right) of purified human cBAF complexes (WT and FF/AA mutant); asterisks indicate non-specific bands. **i.** Remodeling activity of cBAF complexes containing wild-type or FF/AA mutant DPF3a. (Left) Native gel; (right) quantification from four independent experiments. Remodeling activity = ∑(n× Pn) ⁄ (n×∑Pn), Pn is the fraction of remodeled nucleosome populations; data are shown as mean ± s.e.m.

Immunoblot analysis confirmed comparable expression levels of FL and FF/AA DPF3a following reconstitution into Dpf3a KO cells (Fig. 4b). Upon induction of differentiation, ectopic expression of FL DPF3a partially restored formation of elongated multinucleated myotubes, closely resembling WT cultures. In contrast, the FF/AA mutant failed to rescue myotube formation, and mutant-expressing cells remained predominantly mononuclear with markedly impaired fusion capacity (Fig. 4c). These findings demonstrate that HRP2 association is indispensable for the physiological function of DPF3a during myogenic differentiation.

To determine whether the differentiation phenotype reflected restoration of the underlying transcriptional program, we performed RNA-seq analysis following DPF3a reconstitution. Both FL and FF/AA constructs were robustly expressed at the transcript level with high reproducibility across biological replicates (Supplementary Fig. S4a). Re-expression of FL DPF3a partially restored expression of multiple fast myosin heavy chain genes, including Myh3, Myh1, and Myh8, whereas the FF/AA mutant failed to do so (Fig. 4d-e). In contrast, neither FL nor FF/AA DPF3a effectively rescued expression of the Hoxb cluster (Fig. 4f-g). This differential responsiveness suggests that distinct genomic loci exhibit varying dependencies on DPF3a function and temporal chromatin regulation. Whereas the fMyh locus appears capable of partial transcriptional recovery following acute DPF3a restoration, the Hoxb cluster may require sustained or developmentally phased DPF3a activity to establish and maintain appropriate chromatin states. Consistent with earlier analyses, Myod1 expression remained unchanged across all conditions (Supplementary Fig. S4b), further supporting the conclusion that DPF3a acts downstream or independently of the core myogenic lineage determinant.

The inability of the FF/AA mutant to rescue differentiation despite normal expression raised an important mechanistic question: does disruption of HRP2 binding impair the intrinsic remodeling activity of DPF3a-cBAF, or does it selectively compromise chromatin targeting and regulatory function in vivo? To distinguish between these possibilities, we biochemically purified recombinant human cBAF complexes containing either WT or FF/AA mutant DPF3a from HEK293F cells using FLAG affinity purification followed by glycerol gradient sedimentation. Coomassie staining and immunoblot analysis confirmed comparable integrity, purity, and subunit stoichiometry between WT and mutant complexes (Fig. 4h and Supplementary Fig. S4c-d). Importantly, neither preparation contained detectable HRP2 (Supplementary Fig. S4e), allowing direct assessment of intrinsic remodeling activity independent of HRP2 association.

In nucleosome remodeling assays, WT DPF3a-cBAF efficiently mobilized nucleosomes, as evidenced by robust histone octamer sliding along DNA templates. Strikingly, the FF/AA-containing complex displayed remodeling activity comparable to that of WT cBAF, with no significant reduction in catalytic efficiency (Fig. 4i and Supplementary Fig. S4f). These results demonstrate that disruption of HRP2 binding does not intrinsically impair cBAF remodeling capacity. To further validate the specificity of the DPF3a-HRP2 interaction, we purified WT and R527A/R528A (RR/AA) mutant HRP2 proteins, mutations previously shown to disrupt DPF3a association^27^, using tandem GST-FLAG affinity purification from E. coli (Supplementary Fig. S4g-h). Pull-down assays confirmed robust interaction between WT HRP2 and DPF3a-cBAF, whereas the RR/AA mutation abolished binding (Supplementary Fig. S4i).

Collectively, these findings establish HRP2 association as an essential determinant of DPF3a function in vivo. Although the FF/AA mutant retains intact intrinsic remodeling activity, it is incapable of restoring myogenic gene expression or differentiation in primary myoblasts due to defective HRP2 engagement. These observations support a model in which HRP2 functions not as a catalytic cofactor, but as a chromatin-targeting module that links DPF3a-cBAF to H3K36me2/3-marked genomic regions, thereby enabling precise and context-dependent regulation of myogenic transcriptional programs.

### Biochemical reconstitution reveals H3K36me2/3-directed chromatin remodeling by the DPF3a-containing cBAF complex

Having established that disruption of HRP2 binding abolishes DPF3a function in myoblasts without impairing the intrinsic remodeling activity of cBAF, we next sought to define the mechanistic contribution of HRP2 to DPF3a-cBAF regulation. Because HRP2 recognizes H3K36me2/3 through its PWWP domain, we hypothesized that HRP2 confers substrate selectivity by coupling DPF3a-cBAF to H3K36-methylated nucleosomes. To directly test this model, we established a fully reconstituted biochemical system to examine H3K36me2/3-dependent chromatin remodeling by purified DPF3a-cBAF complexes.

Guided by the approximately 226 bp spacing between DPF3a and MyoD peak centers observed in vivo (Fig.3h), a distance corresponding roughly to a single nucleosome plus linker DNA, we designed mononucleosome substrates that recapitulate this chromatin architecture. Recombinant nucleosomes contained asymmetric linker DNA, including a 65 bp linker on one side and a 75 bp linker harboring a Gal4 binding sequence on the other, together with site-specific H3K36me2 or H3K36me3 modifications introduced using the methyl-lysine analog (MLA) strategy^42^ (Fig. 5a-b). This system enabled simultaneous interrogation of histone modification recognition and transcription factor-directed remodeling.

**Figure 5.**
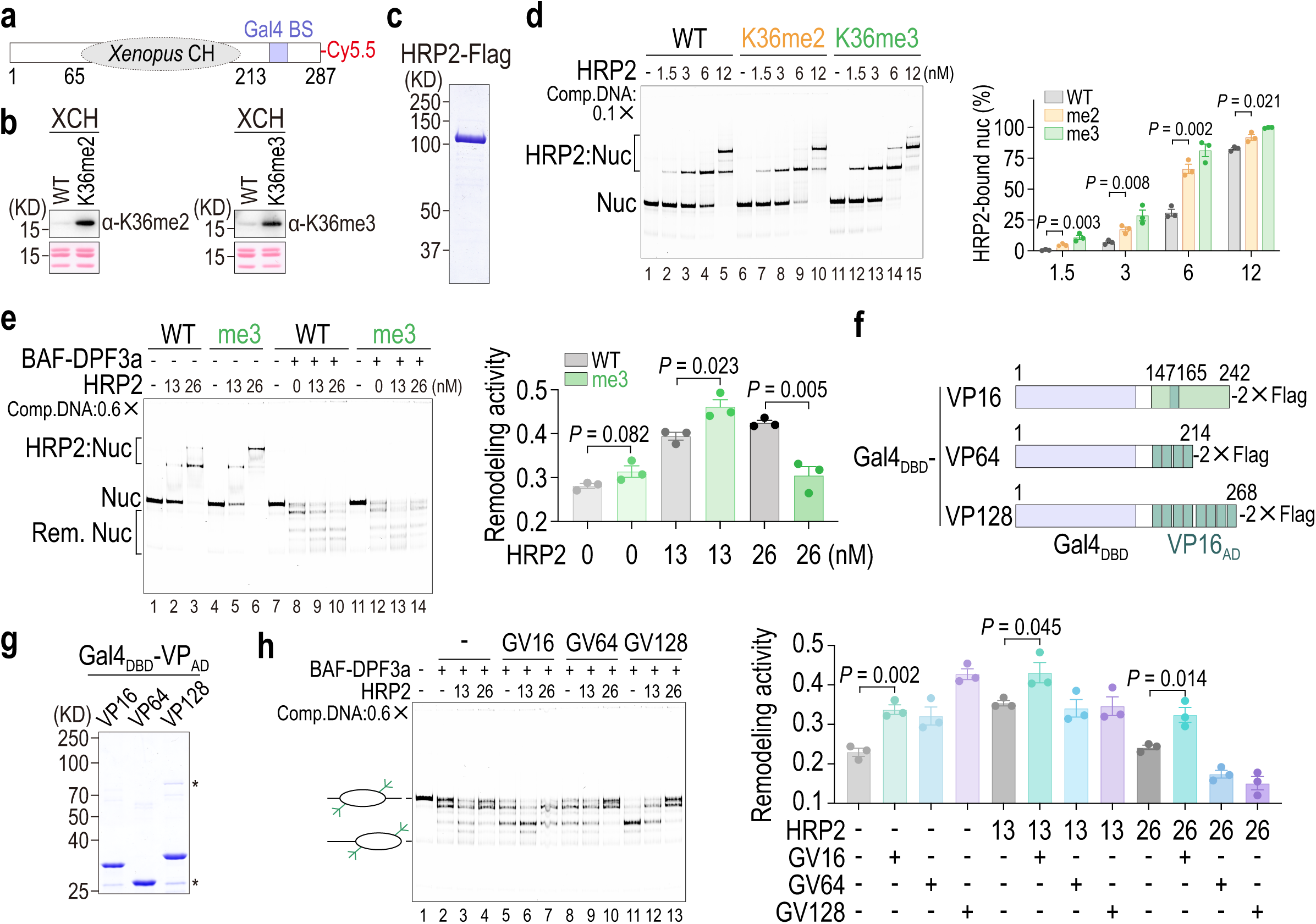
Biochemical reconstitution reveals H3K36me2/3-directed chromatin remodeling by the DPF3a-containing cBAF complex. **a.** Schematic of Cy5.5-labeled reconstituted nucleosome. *Xenopus* core histones (*Xenopus* CH) and the Gal4 binding site (Gal4 BS) are indicated; positions 66-212 correspond to the 601 sequence. **b.** Validation of H3K36me2/3-modified *Xenopus* core histones (XCH) by western blot. **c.** Coomassie blue staining of purified human Flag-tagged HRP2 protein; asterisks indicate non-specific bands. **d.** Electrophoretic mobility shift assay (EMSA) showing binding of HRP2 to reconstituted nucleosomes (WT, H3K36me2 or H3K36me3). (Left) EMSA gel; (right) quantification from three independent experiments. Comp.DNA, competitor DNA. Data are shown as mean ± s.e.m. **e.** Remodeling activity of DPF3a-cBAF on WT or H3K36me3 nucleosomes in the presence of increasing HRP2 (0, 13 and 26 nM). (Left) Native gel; (right) quantification from three independent experiments. Nuc, nucleosomes; Rem. Nuc, remodeled nucleosomes. Data are shown as mean ± s.e.m. **f.** Schematic of Gal4 DNA-binding domain (DBD) fusion proteins: Gal_DBD_-VP16, Gal4_DBD_-VP64 and Gal4_DBD_-VP128. The dark green boxes represent the individual core VP16 motifs. **g.** Coomassie blue staining of purified Gal4_DBD_ fusion proteins shown in **f**; asterisks indicate non-specific bands. **h.** Targeted remodeling activity of DPF3a-cBAF on H3K36me3 nucleosomes using Gal4_DBD_ fusion proteins as recruiters, with increasing HRP2 concentrations (0, 13 and 26 nM). (Left) Native gel; (right) quantification from three independent experiments. GV16, Gal4_DBD_-VP16; GV64, Gal4_DBD_-VP64; GV128, Gal4_DBD_-VP128. Data are shown as mean ± s.e.m.

We first examined whether purified HRP2 directly recognizes modified nucleosomes. Electrophoretic mobility shift assays (EMSA) demonstrated efficient binding of HRP2 to both H3K36me2- and H3K36me3-modified nucleosomes, with modestly higher affinity toward H3K36me3 templates (Fig. 5c-d). Competition assays using unlabeled DNA confirmed the specificity of these interactions (Supplementary Fig. S5a). These findings establish HRP2 as a direct reader of H3K36 methylation capable of linking histone modification recognition to nucleosomal substrate engagement.

We next investigated whether HRP2 modulates DPF3a-cBAF remodeling activity on methylated nucleosomes. In the absence of HRP2, DPF3a-cBAF exhibited comparable remodeling activity on unmodified and H3K36me3-modified nucleosomes (Fig. 5e, lanes 8 and 12). Addition of HRP2 at 13 nM selectively enhanced remodeling of H3K36me3 nucleosomes relative to unmodified templates (Fig. 5e, lanes 9 and 13). Strikingly, increasing HRP2 concentration to 26 nM reversed this effect, suppressing remodeling activity on H3K36me3 nucleosomes below that observed on WT substrates (Fig. 5e, lanes 10 and 14; Supplementary Fig. S5b-d). This biphasic response correlated with distinct HRP2-nucleosome binding behaviors observed by EMSA. At intermediate concentrations, HRP2 preferentially formed a faster-migrating nucleoprotein species associated with enhanced remodeling activity, whereas higher concentrations promoted formation of slower-migrating complexes coincident with remodeling inhibition. These observations suggest that HRP2 exerts concentration-dependent control over DPF3a-cBAF activity by shifting between remodeling-permissive and inhibitory chromatin engagement states.

In vivo, cBAF complexes are frequently recruited to chromatin through interactions with transcription factors and regulatory cofactors. To mimic this mode of recruitment *in vitro*, we introduced Gal4 DNA-binding domain (DBD) fusion proteins carrying activation domains of increasing strength, including VP16, VP64, and VP128^62,63^ (Fig. 5f-g). The engineered Gal4 binding sequence within the linker DNA allowed targeted recruitment of remodeling complexes to modified nucleosomal substrates. Under these conditions, Gal4DBD-VP16 supported efficient HRP2-dependent remodeling of H3K36me3 nucleosomes and reproduced the same biphasic response observed in the untargeted system, with stimulation at intermediate HRP2 levels and inhibition at higher concentrations (Fig. 5h and Supplementary Fig. S5e). Surprisingly, stronger recruitment mediated by Gal4DBD-VP128 reduced remodeling efficiency even at intermediate HRP2 concentrations and further suppressed activity under high-HRP2 conditions. EMSA analyses confirmed stable formation of supershifted complexes containing HRP2, Gal4 fusion proteins, and H3K36me3 nucleosomes (Supplementary Fig. S5f), indicating that reduced remodeling efficiency did not arise from defective chromatin binding, but rather from excessive recruitment strength itself. Thus, productive remodeling requires balanced chromatin engagement, whereas overly stable or excessive recruitment becomes inhibitory.

Collectively, these biochemical reconstitution experiments establish a direct mechanistic link between H3K36 methylation recognition and ATP-dependent chromatin remodeling by the DPF3a-cBAF complex. HRP2 couples DPF3a-cBAF to H3K36me2/3-modified nucleosomes, yet this stimulation occurs only within a defined concentration window, with excessive HRP2 engagement suppressing remodeling activity. Similarly, targeted recruitment enhances remodeling only within an optimal range, whereas excessive recruitment strength becomes counterproductive. These findings reveal that DPF3a-cBAF activity is governed by finely tuned regulatory constraints integrating histone modification recognition, cofactor abundance, and chromatin engagement dynamics, thereby ensuring spatially and contextually restricted remodeling activity within the genome.

### In contrast to canonical remodeling paradigms, DPF3a-BAF exerts limited effects on local chromatin accessibility at functional target loci

The extensive transcriptional perturbations observed upon Dpf3a deletion (Fig. 2) prompted us to examine whether these changes are accompanied by corresponding alterations in chromatin accessibility. We therefore analyzed ATAC-seq profiles from WT and Dpf3a KO myoblasts and systematically compared accessibility dynamics with transcriptional changes. To contextualize these findings within established BAF remodeling paradigms, we reanalyzed published datasets from SMARCE1-deficient cells^61^ using the same computational pipeline. SMARCE1 is a core subunit shared between cBAF and PBAF complexes^64^ and serves as a representative mediator of canonical BAF-dependent chromatin remodeling. This comparison allowed us to distinguish the regulatory behavior of the tissue-specific subunit DPF3a from that of broadly acting core remodeling machinery.

Genome-wide occupancy profiles revealed substantial promoter-proximal localization for both DPF3a and SMARCE1 (Fig. 6a-b and Fig. 6e-f). Genes associated with promoter-bound peaks were subsequently defined as target genes, and their transcriptional responses were evaluated following loss of each factor. Surprisingly, despite widespread transcriptomic alterations in Dpf3a KO cells, only a small fraction of DPF3a promoter-associated target genes exhibited significant expression changes, with 0.7% upregulated and 1.64% downregulated (Fig. 6c). In contrast, SMARCE1 target genes displayed substantially broader transcriptional perturbation, with 8.51% upregulated and 9.57% downregulated upon SMARCE1 depletion (Fig. 6g). These observations suggest that DPF3a does not primarily regulate transcription through direct and widespread promoter-centered remodeling, but instead exerts more selective and spatially restricted control over gene expression.

**Figure 6.**
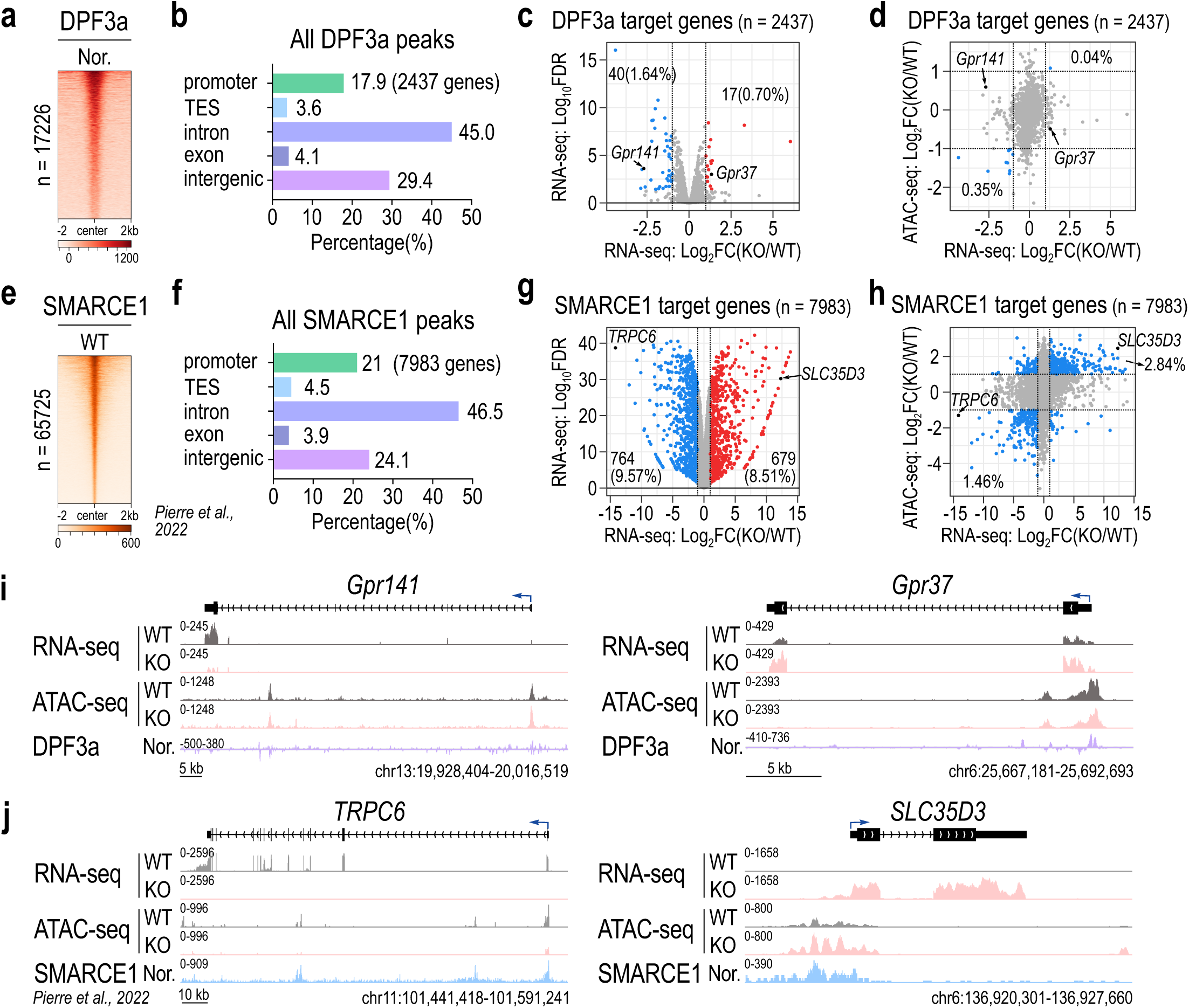
In contrast to canonical remodeling paradigms, DPF3a-BAF exerts limited effects on local chromatin accessibility at functional target loci. **a.** Heatmap of all DPF3a binding regions. ‘Nor.’ indicates normalized DPF3a signal generated by subtracting non-specific signal present in the *Dpf3a* KO myoblasts. **b.** Distribution of DPF3a binding regions by genomic feature (promoter, TES, intron, and intergenic). **c.** Volcano plot depicting the expression profiles of DPF3a target genes identified from promoter-associated binding regions (as defined in **b**). Dashed lines indicate |log_2_FC| =1. **d.** Scatter plot correlating promoter accessibility log_2_FC (KO/WT) with gene expression log_2_FC (KO/WT) for DPF3a target genes with annotated promoters as defined in **b**, with representative genes indicated. Quadrants highlight genes with |log_2_FC| >1 in both assays **e.** Heatmap of published SMARCE1 binding regions **f.** Distribution of SMARCE1 binding regions by genomic feature (promoter, TES, intron, and intergenic) **g.** Volcano plot depicting the expression profiles of SMARCE1 target genes identified from promoter-associated binding regions (as defined in **f**). Dashed lines indicate |log_2_FC| =1. **h.** Scatter plot correlating promoter accessibility log_2_FC (KO/WT) with gene expression log_2_FC (KO/WT) for SMARCE1 target genes with annotated promoters as defined in **b**, with representative genes indicated. Quadrants highlight genes with |log_2_FC| >1 in both assays. **i.** Representative genome browser tracks of DPF3a target genes (as indicated in **c**) that exhibit altered expression but no significant change in promoter accessibility. **j.** Representative genome browser tracks of SMARCE1 target genes (as indicated in **g**) that exhibit both altered expression and a significant change in promoter accessibility.

We next examined whether transcriptional changes at DPF3a target genes correlate with altered promoter accessibility. Remarkably, promoter accessibility and transcriptional output were largely uncoupled in DPF3a-regulated genes. Only 0.04% of target genes exhibited concordant increases in both accessibility and expression, whereas 0.35% displayed concordant decreases (Fig. 6d). By comparison, SMARCE1 target genes showed substantially stronger concordance, with 2.84% exhibiting coupled increases and 1.46% exhibiting coupled decreases in accessibility and transcription (Fig. 6h). These findings indicate that DPF3a-mediated transcriptional regulation operates largely independently of promoter accessibility changes, in marked contrast to the canonical remodeling paradigm exemplified by SMARCE1.

Genome browser analyses of representative loci further illustrated this distinction. At DPF3a target genes such as Gpr141 and Gpr37, clear promoter occupancy by DPF3a was detectable, yet promoter accessibility remained essentially unchanged between WT and Dpf3a KO myoblasts despite significant transcriptional alterations (Fig. 6i). Conversely, canonical SMARCE1 target genes including Trpc6 and Slc35d3 exhibited coordinated changes in both promoter accessibility and gene expression following SMARCE1 loss (Fig. 6j), consistent with conventional accessibility-dependent remodeling mechanisms.

To further evaluate the global relationship between DPF3a-dependent accessibility changes and transcriptional regulation, we classified ATAC-seq peaks into increased, decreased, and unchanged accessibility groups in both DPF3a and SMARCE1 datasets (Supplementary Fig. S6a, c). Genomic annotation revealed that accessibility changes associated with DPF3a loss were enriched primarily within intergenic and intronic regions, whereas accessibility-stable regions were predominantly promoter-associated (Supplementary Fig. S6b). In contrast, SMARCE1-dependent accessibility changes displayed distinct genomic distributions, with decreased-accessibility regions preferentially enriched within intronic chromatin (Supplementary Fig. S6d). These divergent patterns suggest that loss of distinct BAF subunits produces fundamentally different chromatin responses.

Cross-analysis of promoter accessibility and transcriptional output further reinforced the uncoupling between accessibility and gene regulation in the DPF3a system. Among promoter-associated regions, concordant increases in accessibility and expression were observed in only 1.19% of DPF3a-regulated genes, compared with 2.80% for SMARCE1, whereas concordant decreases occurred in 1.05% and 2.61% of genes, respectively (Supplementary Fig. S6e-f). Thus, unlike canonical BAF remodeling factors, DPF3a exerts transcriptional effects with remarkably limited dependence on local promoter accessibility changes.

Collectively, these findings indicate that DPF3a-cBAF operates through a regulatory mechanism fundamentally distinct from classical accessibility-centered chromatin remodeling. Notably, motif enrichment analysis of DPF3a-regulated accessible regions revealed strong enrichment of CTCF-binding motifs (Supplementary Fig. S6g), implicating DPF3a-associated chromatin regions in higher-order genome organization rather than direct promoter remodeling. These observations raised the possibility that DPF3a regulates transcription through modulation of long-range chromatin interactions.

### DPF3a promotes long-range chromatin looping to regulate myogenic gene expression

The pronounced uncoupling between transcriptional output and promoter accessibility following Dpf3a deletion (Fig. 6), together with the enrichment of CTCF motifs at DPF3a-regulated chromatin regions (Supplementary Fig. S6g), suggested that DPF3a may influence gene expression through higher-order chromatin architecture rather than conventional promoter opening mechanisms. To directly examine this possibility, we performed CTCF ChIP-seq in WT and Dpf3a KO primary myoblasts.

Principal component analysis of CTCF occupancy profiles revealed clear segregation between WT and Dpf3a KO samples (Fig. 7a), indicating that DPF3a deficiency broadly alters the chromatin binding landscape of CTCF. We next examined CTCF occupancy across the three classes of DPF3a-associated ATAC-seq regions defined previously (up, down, and unchanged accessibility; Supplementary Fig. S6a). Heatmap analyses demonstrated that DPF3a loss produced coordinated alterations in CTCF occupancy at both accessibility-increased and accessibility-decreased regions, which were themselves characterized by focal H3K36me2 depressions and prominent MyoD enrichment (Fig. 7b). Meta-profile and violin plot analyses further revealed that accessibility changes were accompanied by concordant shifts in CTCF and H3K36me2 signals (Fig. 7c-d). Regions exhibiting increased accessibility in Dpf3a KO cells displayed relatively low DPF3a occupancy together with elevated CTCF and H3K36me2 signals, whereas regions with decreased accessibility showed the opposite pattern, including enrichment of DPF3a and reduced CTCF and H3K36me2 levels.

**Figure 7.**
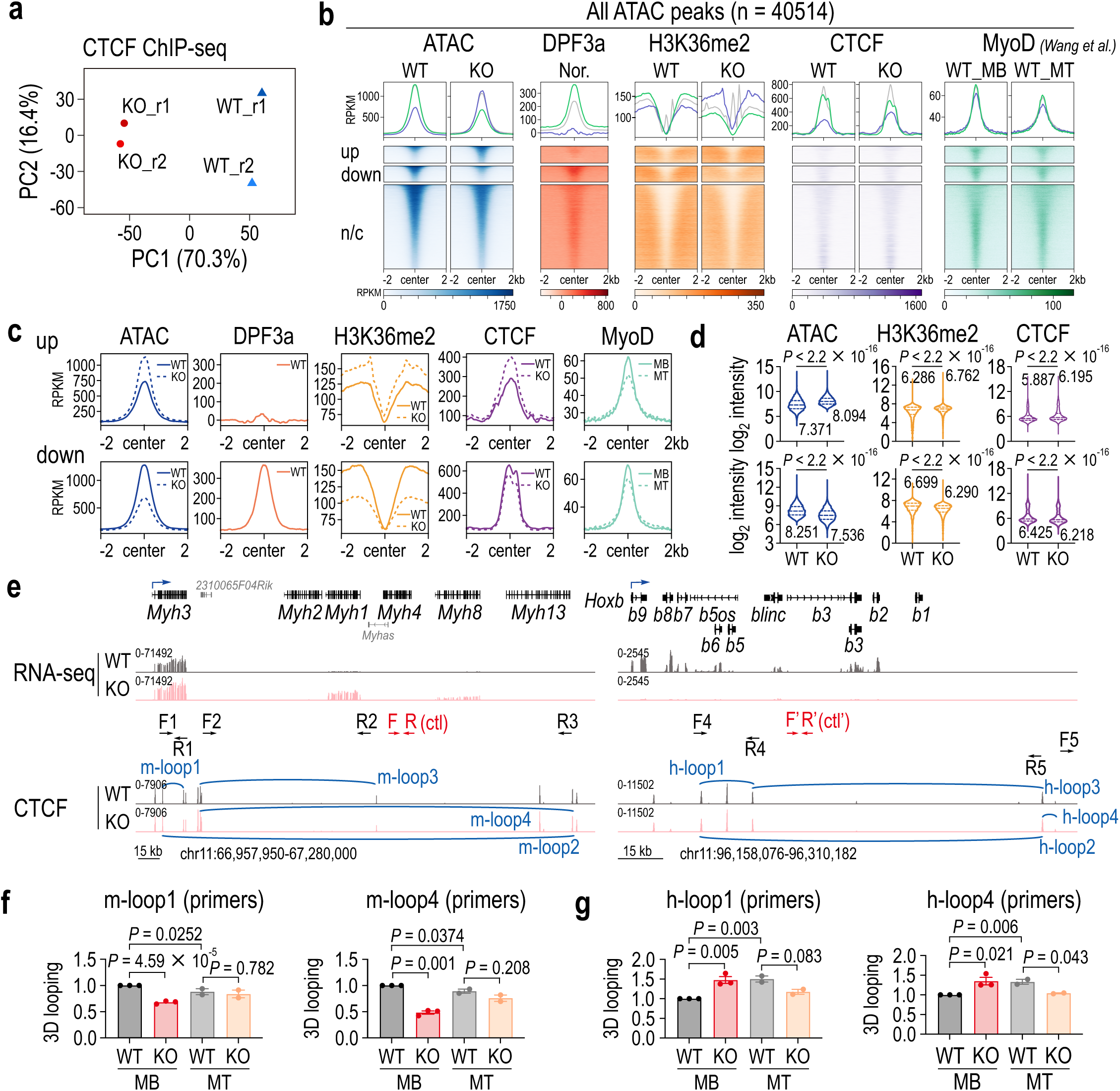
DPF3a promotes long-range chromatin looping to regulate myogenic gene expression. **a.** Principal component analysis (PCA) of CTCF ChIP-seq signals at all binding regions in myoblast. **b.** Heatmap showing CTCF binding patterns at all ATAC-seq peaks, accompanied by normalized PF3a, H3K36me2 and MyoD signals. ‘Nor.’ indicates normalized DPF3a signal generated by subtracting non-specific signal present in the *Dpf3a* KO myoblasts; MB, myoblast; MT, myotube. **c.** Meta-plot showing the distribution of epigenetic signals in “up” and “down” regions. MB, myoblast; MT, myotube. **d.** Violin plot showing the distribution of epigenetic signals in C1 and C2 regions; the numbers above the violins indicate the mean value. **e.** Schematic showing the location of primers used for 3D chromatin interaction detection at the *fMyh* and *Hoxb* loci. Primers used for detection of each loop are as indicated; ctl and ctl’ primers serve as internal controls for qPCR. **f.** qPCR quantification of 3D looping interactions of m-loop1 and m-loop4 at the *fMyh* locus from three independent experiments. Data are shown as mean ± s.e.m. **g.** qPCR quantification of 3D looping interactions of h-loop1 and h-loop4 at the *Hoxb* locus from three independent experiments. Data are shown as mean ± s.e.m.

To quantitatively assess these relationships, we performed correlation analyses between ATAC-seq intensity and chromatin-associated factors within accessibility-altered regions (Supplementary Fig. S7a). Chromatin accessibility exhibited a weak but consistent inverse correlation with H3K36me2 levels in both accessibility-increased and accessibility-decreased regions, whereas accessibility positively correlated with DPF3a, CTCF, and MyoD occupancy. Notably, MyoD displayed strong focal enrichment at both classes of regulatory regions and showed higher occupancy in myoblasts than in differentiated myotubes (Fig. 7c-d and Supplementary Fig. S7a), consistent with recent evidence implicating MyoD as an organizer of myoblast chromatin architecture^65^. Collectively, these findings suggest that DPF3a participates in coordinated regulatory assemblies involving CTCF, MyoD, and H3K36me2-associated chromatin environments at putative architectural elements.

To directly determine whether DPF3a regulates chromatin looping, we focused on the fast myosin heavy chain (fMyh) and Hoxb loci, two genomic regions that exhibited opposing transcriptional responses following Dpf3a deletion (Fig. 2g) and are known to undergo dynamic conformational remodeling during differentiation^35,36^. Guided by CTCF occupancy patterns, we designed locus-specific chromosome conformation capture (3C) assays to quantify point-to-point chromatin interactions across these regions (Fig. 7e and Supplementary Fig. S7b), with assay specificity validated by multiple control experiments (Supplementary Fig. S7c-d).

At the fMyh locus, we identified four prominent CTCF-associated chromatin loops (m-loop1-4) (Fig. 7e). Consistent with our transcriptomic analyses, both RT-qPCR and RNA-seq confirmed precocious activation of fMyh genes in Dpf3a KO myoblasts (Supplementary Fig. S7e-f). Strikingly, 3C analyses revealed that interactions corresponding to m-loop1 and m-loop4 were significantly weakened in Dpf3a-deficient myoblasts relative to WT controls (Fig. 7f and Supplementary Fig. S7g). As an additional validation, we examined loop dynamics during normal differentiation. In WT myotubes, interactions at m-loop1 and m-loop4 were markedly diminished compared with WT myoblasts, consistent with previous Hi-C studies demonstrating dissolution of the fMyh chromatin interaction domain during myogenic differentiation^35^. These results establish that DPF3a is required to maintain specific chromatin loops at the fMyh locus and further suggest that disruption of these architectural interactions is associated with premature activation of fast myosin genes.

We next examined chromatin architecture at the Hoxb locus using four corresponding interaction pairs (h-loop1-4) (Fig. 7e). In contrast to the fMyh region, Dpf3a deficiency resulted in significantly increased interactions at h-loop1 and h-loop4 (Fig. 7g and Supplementary Fig. S7h), correlating with the widespread repression of Hoxb genes observed in Dpf3a KO cells (Fig. 2g). Analysis of differentiated myotubes further revealed an overall reduction in looping interactions at both fMyh and Hoxb loci, with particularly pronounced decreases at the Hoxb region. These observations likely reflect the cumulative architectural and transcriptional dysregulation that emerges as differentiation proceeds in the absence of DPF3a. Importantly, the opposing loop alterations observed at the two loci—loop loss at fMyh and loop gain at Hoxb—indicate that DPF3a does not simply function as a global promoter or suppressor of chromatin looping. Rather, DPF3a appears to maintain locus-specific three-dimensional chromatin architectures compatible with appropriate transcriptional states.

Together, these findings demonstrate that DPF3a regulates gene expression through modulation of long-range chromatin looping. DPF3a deficiency induces distinct architectural perturbations at different genomic loci, with loop destabilization at fMyh associated with transcriptional activation and loop strengthening at Hoxb associated with transcriptional repression. These opposing outcomes suggest that DPF3a functions as a context-dependent architectural regulator that preserves the appropriate chromatin topology required for lineage-specific gene control. Given that DPF3a localizes to focal depressions within H3K36me2 domains and exhibits HRP2-dependent remodeling activity (Fig. 3-5), our findings support a model in which DPF3a-cBAF couples histone modification recognition to selective nucleosome remodeling at chromatin architectural elements, thereby facilitating or stabilizing CTCF-associated long-range interactions. This mechanism operates largely independently of promoter accessibility and provides a direct conceptual link between histone modification-guided chromatin remodeling and higher-order genome organization during myogenic differentiation.

## Discussion

In this study, we identify DPF3a as a tissue-specific regulator that links histone modification sensing to higher-order genome organization during skeletal muscle differentiation and regeneration. Although ATP-dependent chromatin remodelers are classically viewed as regulators of local chromatin accessibility through nucleosome repositioning ^2,3^, our findings support a distinct regulatory paradigm in which a specialized cBAF complex controls transcription predominantly through long-range chromatin architecture. Mechanistically, DPF3a cooperates with the H3K36me2/3 reader HRP2, preferentially localizes to focal depressions embedded within broad H3K36me2 domains, and regulates CTCF-associated chromatin looping at key myogenic loci. Together, these findings establish a direct mechanistic connection between histone modification-guided chromatin remodeling and three-dimensional genome organization in lineage-specific gene regulation.

A central conceptual advance of this work is the identification of a distinctive chromatin configuration that guides DPF3a localization. Rather than occupying regions of maximal H3K36me2 enrichment, DPF3a preferentially localizes to focal depressions nested within broad H3K36me2 plateaus. This organization differs fundamentally from conventional models in which chromatin-associated reader proteins simply track the abundance of their cognate histone modifications ^66,67^. Our data instead support a hierarchical targeting model in which broad H3K36me2 domains establish permissive chromatin territories, while lineage-determining transcription factors such as MyoD refine the precise recruitment sites of DPF3a-cBAF. The sharp co-enrichment of MyoD at these regions, together with the characteristic ∼226 bp spatial offset between DPF3a and MyoD peak centers, further suggests a highly ordered nucleosomal arrangement. One possibility is that DPF3a-cBAF remodels nucleosomes adjacent to MyoD-bound DNA rather than directly overlapping transcription factor occupancy, thereby permitting remodeling without destabilizing lineage-specific transcription factor binding. More broadly, this configuration may represent a general strategy by which chromatin remodelers achieve positional specificity within expansive histone modification landscapes while preserving local transcription factor engagement.

Our biochemical reconstitution experiments further reveal that HRP2 functions as a regulatory module that couples H3K36 methylation recognition to DPF3a-cBAF remodeling activity. Although HRP2 association is indispensable for DPF3a function in vivo, disruption of this interaction does not impair intrinsic remodeling activity of the cBAF complex itself, indicating that HRP2 primarily acts through substrate engagement and chromatin targeting rather than catalytic activation. Importantly, remodeling stimulation occurred only within a restricted concentration range of HRP2, with higher concentrations becoming inhibitory. This biphasic behavior correlated with distinct nucleoprotein species observed in EMSA assays, suggesting that alternative HRP2-nucleosome binding states differentially influence remodeling competency. Such a mechanism may provide an intrinsic buffering system that constrains remodeling activity within appropriate chromatin contexts and prevents excessive or ectopic nucleosome mobilization. Similarly, our observation that moderate recruitment strength promotes remodeling more effectively than excessive recruitment indicates that chromatin remodeling efficiency is governed by a non-linear relationship between targeting avidity and catalytic output. These findings raise the possibility that productive remodeling requires a dynamic and transient engagement state rather than maximal chromatin occupancy.

Perhaps the most unexpected finding of this study is the pronounced uncoupling between promoter accessibility and transcriptional output in the DPF3a system. Despite robust transcriptional dysregulation following Dpf3a deletion, accessibility changes at promoters of directly bound genes were remarkably limited. This behavior contrasts sharply with canonical BAF remodeling paradigms exemplified by the core subunit SMARCE1, where transcriptional changes closely parallel promoter accessibility alterations. Instead, DPF3a-dependent accessible regions were enriched for CTCF motifs and associated with distal regulatory chromatin, suggesting that DPF3a primarily functions through architectural regulation rather than promoter opening. These observations challenge the prevailing assumption that ATP-dependent remodelers regulate transcription principally through accessibility modulation and instead indicate that remodelers can exert substantial transcriptional control while leaving local promoter accessibility largely unchanged.

Our chromatin conformation analyses provide direct evidence supporting this architectural mechanism. At both the fMyh and Hoxb loci, DPF3a loss altered long-range chromatin interactions in a locus-dependent manner that closely correlated with transcriptional changes. Notably, the directionality of loop perturbation differed between loci: loop destabilization at the fMyh locus accompanied precocious transcriptional activation, whereas increased looping at the Hoxb locus correlated with transcriptional repression. These findings indicate that DPF3a does not function as a simple promoter or suppressor of chromatin looping. Rather, DPF3a appears to preserve locus-specific chromatin topologies compatible with appropriate transcriptional states. Such context dependency likely reflects differences in local CTCF architecture, chromatin environment, transcription factor occupancy, and associated cofactors across distinct genomic regions. Importantly, the fMyh and Hoxb loci likely represent broader classes of DPF3a-regulated chromatin domains rather than isolated examples, suggesting that architectural regulation constitutes a widespread mechanism underlying DPF3a-dependent myogenic transcription programs.

Our findings also contribute to an emerging framework linking chromatin remodeling complexes to genome architecture through mechanistically distinct pathways. In a recent study, Sun et al. demonstrated that the NuRD complex stabilizes CTCF occupancy and promotes TET-dependent DNA demethylation at CTCF-binding sites, thereby maintaining local hypomethylation and higher-order chromatin organization during stem cell differentiation^68^. Independent studies have additionally established NuRD as a key regulator of striated muscle identity and lineage fidelity^69,70^. Together with our work, these findings suggest that multiple chromatin remodeling systems converge on the regulation of genome architecture through distinct molecular mechanisms. Whereas NuRD appears to maintain permissive chromatin environments at CTCF sites through DNA methylation control^68^, DPF3a-cBAF instead links H3K36 methylation sensing to nucleosome remodeling at distal architectural elements without broadly altering promoter accessibility. These observations collectively support the emerging concept that chromatin remodelers contribute to three-dimensional genome regulation through specialized and context-dependent mechanisms extending beyond canonical accessibility modulation.

Several limitations should be considered when interpreting our findings. First, although our biochemical reconstitution assays establish direct H3K36me2/3-dependent remodeling by DPF3a-cBAF, these systems necessarily simplify the complexity of native chromatin environments, including the contributions of additional chromatin-associated factors and higher-order chromatin topology. Second, our analyses of transcriptional and accessibility uncoupling were based primarily on population-level genomic assays; future single-cell multi-omic approaches may reveal additional layers of heterogeneity and temporal regulation during differentiation. Third, while targeted 3C analyses demonstrated locus-specific looping defects at representative myogenic regions, genome-wide chromatin conformation profiling will be required to define the full extent of DPF3a-dependent architectural regulation across the genome.

Based on these findings, we propose a working model in which DPF3a-cBAF is recruited to focal depressions within broad H3K36me2 domains through cooperative interactions involving HRP2 and lineage-specific transcription factors such as MyoD. At these sites, DPF3a-cBAF remodels nucleosomes in a manner that facilitates or stabilizes CTCF-associated long-range chromatin interactions, thereby coordinating transcriptional programs required for myoblast differentiation and muscle regeneration. In this framework, DPF3a functions not as a general accessibility factor, but as a specialized architectural regulator that integrates histone modification landscapes with higher-order genome folding. More broadly, our study expands the conceptual framework of chromatin remodeling by demonstrating that ATP-dependent remodelers can regulate transcription through modulation of three-dimensional genome organization independently of local promoter accessibility changes. Given the extensive involvement of BAF complex mutations in developmental disorders and cancer^71^, it will be important to determine whether dysregulation of DPF3a-dependent architectural remodeling contributes to muscle pathologies and other lineage-associated diseases.

## Data availability

The data that support this study are available from the corresponding authors upon reasonable request. The raw sequence data reported in this paper have been deposited under accession numbers CRA037139 in the Genome Sequence Archive (GSA)^72^ in National Genomics Data Center.

Published ATAC-seq and RNA-seq for SMARCE1 are obtained from GEO under accession number GSE174360; ChIP-seq for MyoD, H3K4me1, H3K4me3, H3K27ac and H3K27me3 are obtained from GSA under accession number CRA002490; ChIP-seq for H3K36me2 and H3K36me3 in mESCs are obtained from GEO under accession number GSE118785.

Custom scripts used for data processing and analysis have been deposited in Zenodo (https://doi.org/10.5281/zenodo.20151856). All software tools and packages employed in this study are publicly available, and their respective versions are specified in the Methods section. Processed bigWig files have additionally been deposited in Zenodo (https://doi.org/10.5281/zenodo.20460044).

## Supplementary data

Supplementary data is available.

## Acknowledgements

We thank the Experimental Flow Cytometry Laboratory of the Core Facility of Basic Medical Sciences and the Animal Science Laboratory at Shanghai Jiao Tong University School of Medicine for technical assistance. We also thank Dr. Qing Zhong (Shanghai Jiao Tong University School of Medicine) for generously providing the HEK293F cell line.

## Author contributions

B.L. and M.Y. designed the study with contributions from the other authors; M.Y. and Y.L. performed CTX model; M.Y. conducted myoblast differentiation with the help of Y. C.; M.Y. performed adenovirus infection and 3C assay with the help of P.H.; M.Y., J.X. and J.W. performed remodeling assay; M.Y., Q.Z. and Y.P. conducted genomic assays; M.Y. and J.X. performed the bioinformatic analysis; M.Y., Y.Y. and M.P. constructed the plasmids; M.Y. purified proteins with the help of M.H., M.L. and Y.F.; B.L. and M.Y. wrote the manuscript; B.L. supervised the project.

## Funding

This work was supported by the National Key R&D Program of China (2021YFA1300100 to B.L.), the National Natural Science Foundation of China (32030019, 31872817 to B.L., 32500488 to J.X., 81901284 to YH.Y.), Natural Science Foundation of Shanghai (25ZR1402294 to Y.P., 25ZR1402295 to J.X.) and China Postdoctoral Science Foundation (2025M772709 to Y.P., 2025M772802 to J.X.).

## Conflict of interests

The authors declare no competing interests.

## Notes

### Competing Interest Statement

The authors have declared no competing interest.

