## Supplementary Data for "DPF3a couples H3K36me2-dependent chromatin remodeling to genome architecture during myogenesis"

#### Supplementary Figures

**Supplementary Figure S1. Isolation of *Dpf3a* knockout muscle stem cells.**

**a.** Expression levels of *DPF1*, *DPF2* and *DPF3* in normal human tissues (data from GTEx database).

**b.** Validation of *Dpf3a* knockout mice in myoblasts derived from muscle tissue by PCR genotyping.

**c.** Schematic of isolation of primary muscle stem cells (MuSCs) by fluorescence-activated cell sorting (FACS).

**d.** Fluorescence-activated cell sorting of mouse MuSCs. After excluding debris and doublets, $VACM1^{+}$ $Sca1^{-}CD31^{-}CD45^{-}$ MuSCs were sorted.

**Supplementary Figure S2. DPF3a acts as a central transcriptional co-regulator of the myogenic gene expression program.**

**a.** Genome browser tracks of RNA-seq signals at the *Myod1* locus in WT and *Dpf3a* KO myoblasts across all time points.

**b.** (Left) FPKM of *Myod1* during myoblast differentiation (days 0, 0.25, 0.5, 1, 2 and 3) in WT and *Dpf3a* KO cells. (Right) RT-qPCR quantification from three independent experiments of *Myod1* at days 0, 1, 2 and 3; data are shown as mean ± s.e.m.

**c.** (Top) Scaled expression (Scaled exp.) and (bottom) average expression (Ave. exp.) dynamics of the R1, R3 and R5 cluster genes identified in **Fig. 2d**

**d.** Gene Ontology (GO) analysis (Biological Processes) of the R3 cluster genes shown in **Fig. 2d.**

**e.** FPKM of *Myh1*, *Myh4* and *Myh8* during myoblast differentiation (days 0, 0.25, 0.5, 1, 2 and 3) in WT and *Dpf3a* KO cells.

**f.** Genome browser tracks of RNA-seq signals at the *Dpf3* locus. (Left) exons 10-12; (right) exons 8-9.

**g.** RT-qPCR quantification of *Dpf3a* at days 0, 1, 2 and 3 from three independent experiments.

**h.** Western blot analysis of DPF3a protein expression dynamics during myoblast differentiation (days 0, 1 and 2) in WT and *Dpf3a* KO cells.

**i.** RT-qPCR quantification of *Dpf3b* at days 0, 1, 2 and 3 from three independent experiments; data are shown as mean ± s.e.m.

**j.** Volcano plot of differentially expressed genes (DEGs) between WT and *Dpf3a* KO myoblasts at day 0.

**k.** Top 5 enriched GO Biological Processes terms for up-regulated and down-regulated DEGs at day 0.

**Supplementary Figure S3. DPF3a preferentially occupies focal depressions embedded within broad H3K36me2 chromatin domains.**

**a.** Heatmap of Pearson correlation coefficients for ATAC-seq, H3K36me2 and H3K36me3 ChIP-seq in WT and *Dpf3a* KO myoblasts.

**b.** Heatmap showing H3K36me3 and normalized H3 signals across all gene regions (-2 kb to TSS to TES to +2kb) in WT myoblasts.

**c.** Genome browser tracks of RNA-seq and H3K36me2 signals in WT myoblasts (this study) and H3K36me2/3 signals in mESCs at the chr12:86,679,578-87,550,745 region (top) and the *Myod1* locus (bottom).

**d.** ChromHMM emission models generated using epigenomic data. Abrrev., abbreviation; Cov., coverage; Annot., annotation.

**e.** Violin plot showing the distribution of epigenetic signals in G1 and G2 DPF3a-binding regions; the numbers above the violins indicate the mean value.

**f.** Representative genome browser tracks at the *fMyh* genomic locus in WT and *Dpf3a* KO myoblasts. Top, the entire *fMyh* locus; bottom, zoom-in view of the region indicated by the box on the top. NFR, nucleosome free region.

**g.** Meta-plot showing the distribution of ATAC signals in overlapping (top) and offset peaks (bottom) as defined in **Fig. 2g**.

**h.** Distribution of overlapping (top) and offset peaks (bottom) by genomic feature (promoter, TES, intron, and intergenic).

**i.** Volcano plot of overlapping (top) and offset peaks (bottom) identified from promoter-associated regions (as defined in **h**). Dashed lines indicate $|\log_2 FC| = 1$. Prom., promoter.

**Supplementary Figure S4. Association with the H3K36me2/3 reader HRP2 is essential for DPF3a function in vivo.**

**a.** Genome browser tracks of RNA-seq signals at chr12:83,316,283-83,316,309 within *Dpf3* exon 9 in WT (empty vector; EV) and KO (EV, FL and FF/AA) myoblasts.

**b.** Genome browser tracks of RNA-seq signals at the *Myod1* locus in WT (EV) and KO (EV, FL and FF/AA) myoblasts.

**c.** Schematic of the purification workflow for human BAF complex containing WT or FF/AA mutant DPF3a.

**d.** Silver staining of fractions from glycerol gradient centrifugation of BAF-DPF3a complex.

**e.** Western blot detection of HRP2 in purified BAF-DPF3a complex (WT and FF/AA); whole cell extract (WCE) and HRP2-Flag protein serve as positive controls; asterisks indicate non-specific bands.

**f.** Schematic of Cy5.5-labeled reconstituted nucleosome. The 65N75 design indicates the 601 sequence flanked by 65 bp (5') and 75 bp (3') linkers; *Xenopus* CH, *Xenopus* core histones.

**g.** Schematic of human HRP2 WT and RR/AA mutant (DPF3a-binding defective); RR/AA, R527A/R528A double mutation in HRP2 that disrupts interaction with HRP2, showed as red lines.

**h.** Coomassie blue staining of purified HRP2 proteins (WT and RR/AA mutant); asterisks indicate non-specific bands.

**i.** GST pull-down of BAF-DPF3a complex using GST-HRP2 (WT or RR/AA). (Left) Western blot; (right) quantification of pull-down efficiency from three independent replicates; data are shown as mean ± s.e.m.

**Supplementary Figure S5. Biochemical reconstitution reveals H3K36me2/3-directed chromatin remodeling by the DPF3a-containing cBAF complex.**

**a.** EMSA showing binding of HRP2 to H3K36me3 nucleosomes in the presence of increasing competitor DNA (unlabeled specific oligonucleotide). Comp.DNA, competitor DNA; Nuc, nucleosome.

**b.** Schematic of the remodeling activity assay for BAF-DPF3a in the presence of HRP2.

**c.** Remodeling activity of BAF-DPF3a (0, 0.6 and 1.3 nM) on H3K36me3 nucleosomes with increasing concentrations of HRP2 (0, 13 and 26 nM).

**d.** Quantification of the remodeling activity shown in **c** from three independent experiments; data are shown as mean ± s.e.m.

**e.** Schematic of the targeted remodeling activity assay for BAF-DPF3a using $Gal4_{DBD}$ fusion proteins as recruiters.

**f.** EMSA showing the supershifted complexes formed by HRP2, $Gal4_{DBD}$ fusion proteins and H3K36me3 nucleosomes. GV16, $Gal4_{DBD}$-VP16; GV64, $Gal4_{DBD}$-VP64; GV128, $Gal4_{DBD}$-VP128. Data are shown as mean ± s.e.m.

**Supplementary Figure S6. Comparison of all altered accessible regions upon *Dpf3a* knockout and *SMARCE1* deficient.**

**a.** Heatmap showing ATAC-seq accessibility dynamics in WT and *Dpf3a* KO myoblasts; n/c, nonchanged.

**b.** Genomic annotation of "up", "down" and "nonchanged (n/c)" clusters for DPF3a as defined in **a**.

**c.** Heatmap showing ATAC-seq accessibility dynamics in WT and *SMARCE1* KO AC7 cells; n/c, nonchanged; AC7, human SMARCE1-deficient arachnoid cells.

**d.** Genomic annotation of "up", "down" and "n/c" clusters for SMARCE1 as defined in **c**. n/c, nonchanged.

**e.** Distribution of annotated changed peaks (up and down) categorized by genomic feature (promoter, TES, intron and intergenic) for DPF3a-regulated ATAC-peaks (this study) and SMARCE1-regulated ATAC-peaks.

**f.** Scatter plots correlating promoter accessibility $\log_2$FC (KO/WT) with gene expression $\log_2$FC (KO/WT) for all genes with annotated promoters as defined in **e**, with representative genes indicated. (Left) This study; (right) published SMARCE1 data; Quadrants highlight genes with $|\log_2FC| > 1$ in both assays (percentages shown). ATAC_prom, changed ATAC-peaks annotated as promoters in **e**.

**g.** Motif enrichment analysis of DPF3a up- and down-regulated differentially accessible peaks (as defined in **a**).

**Supplementary Figure S7. DPF3a promotes long-range chromatin looping to regulate myogenic gene expression.**

**a.** Scatterplot showing correlation levels between ATAC-seq signal and DPF3a, H3K36me2, CTCF and or MyoD at ATAC signal "up" and "down" regions (defined in Fig. 7b). Spearman correlation coefficient (R) and *P* value are displayed.

**b.** Schematic of the chromosome conformation capture (3C) assay used to analyze chromatin conformation changes during differentiation from primary myoblasts to myotubes.

**c.** Agarose gel showing ligated 3C DNA templates; the minus T4 ligase condition serves as a negative control.

**d.** PCR analysis of 3C DNA templates from WT and KO cells. Lanes 1-8: control primers at the *fMyh* locus; lanes 9-16: control primers at the *Hoxb* locus.

**e.** Expression levels of *Myh3* in WT and *Dpf3a* KO myoblasts as assessed by RT-qPCR from three independent experiments; data are shown as mean ± s.e.m.

**f**. Expression levels of *Myh3*, *Myh1*, *Myh4* and *Myh8* in WT and *Dpf3a* KO myoblasts as assessed by RNA-seq from two biological replicates.

**g.** qPCR quantification of 3D looping interactions of m-loop2 and m-loop3 at the *fMyh* locus from three independent experiments. Data are shown as mean ± s.e.m.

**h.** qPCR quantification of 3D looping interactions of h-loop2 and h-loop3 at the *Hoxb* locus from three independent experiments. Data are shown as mean ± s.e.m.

### Figure S1

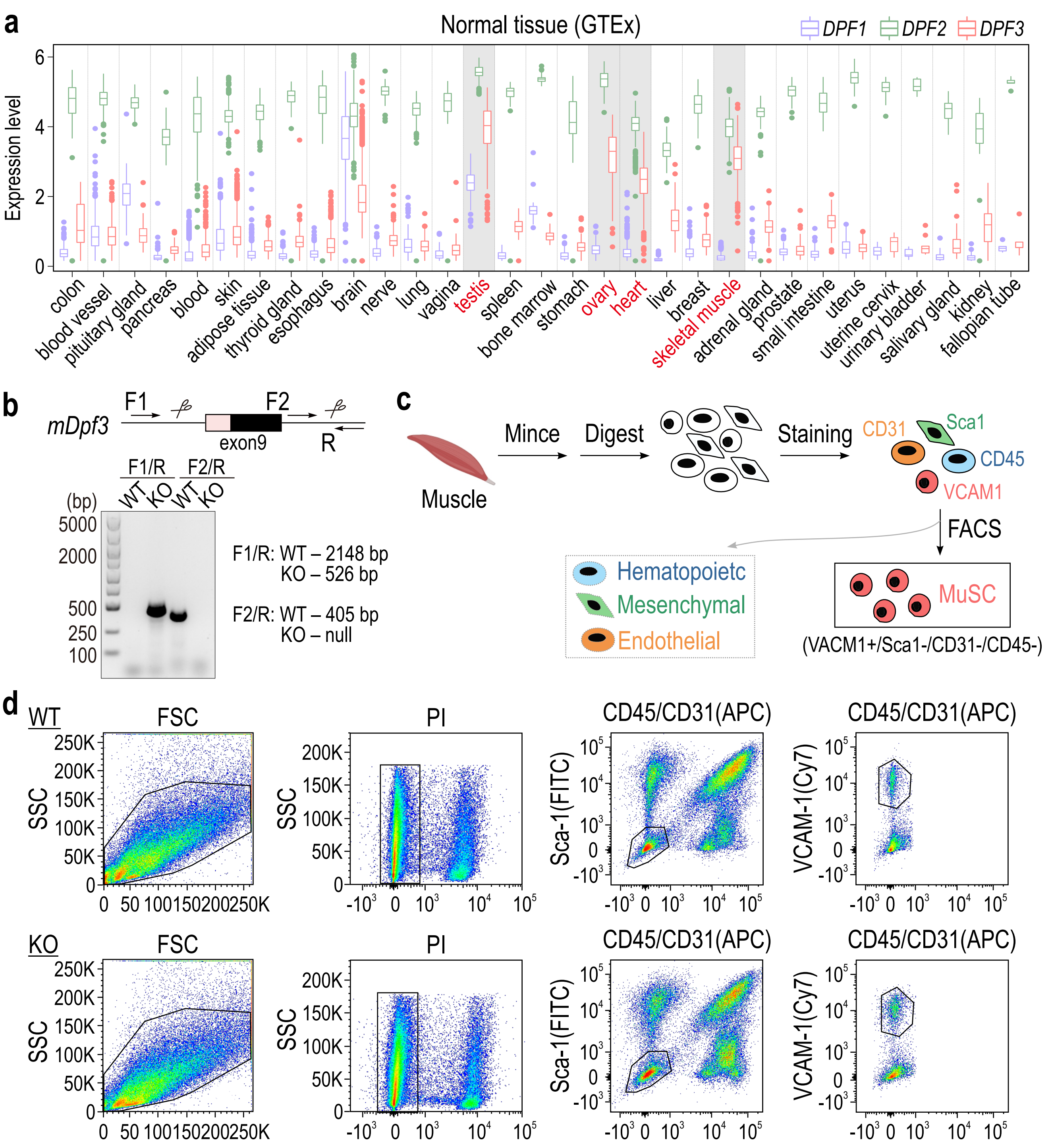
a
Normal tissue (GTEx)
DPF1 DPF2 DPF3
Expression level
colon
blood vessel
pituitary gland
pancreas
blood
skin
adipose tissue
thyroid gland
esophagus
brain
nerve
lung
vagina
testis
spleen
bone marrow
stomach
ovary
heart
liver
breast
skeletal muscle
adrenal gland
prostate
small intestine
uterus
uterine cervix
urinary bladder
salivary gland
kidney
fallopian tube
b
mDpf3
F1
F2
exon9
R
F1/R
F2/R
WT KO WT KO
(bp)
5000
2000
500
250
100
F1/R: WT – 2148 bp
KO – 526 bp
F2/R: WT – 405 bp
KO – null
c
Muscle
Mince
Digest
Staining
CD31
Sca1
CD45
VCAM1
FACS
Hematopoietc
Mesenchymal
Endothelial
MuSC
(VACM1+/Sca1-/CD31-/CD45-)
d
WT
FSC
SSC
PI
CD45/CD31(APC)
Sca-1(FITC)
VCAM-1(Cy7)
KO

### Figure S2

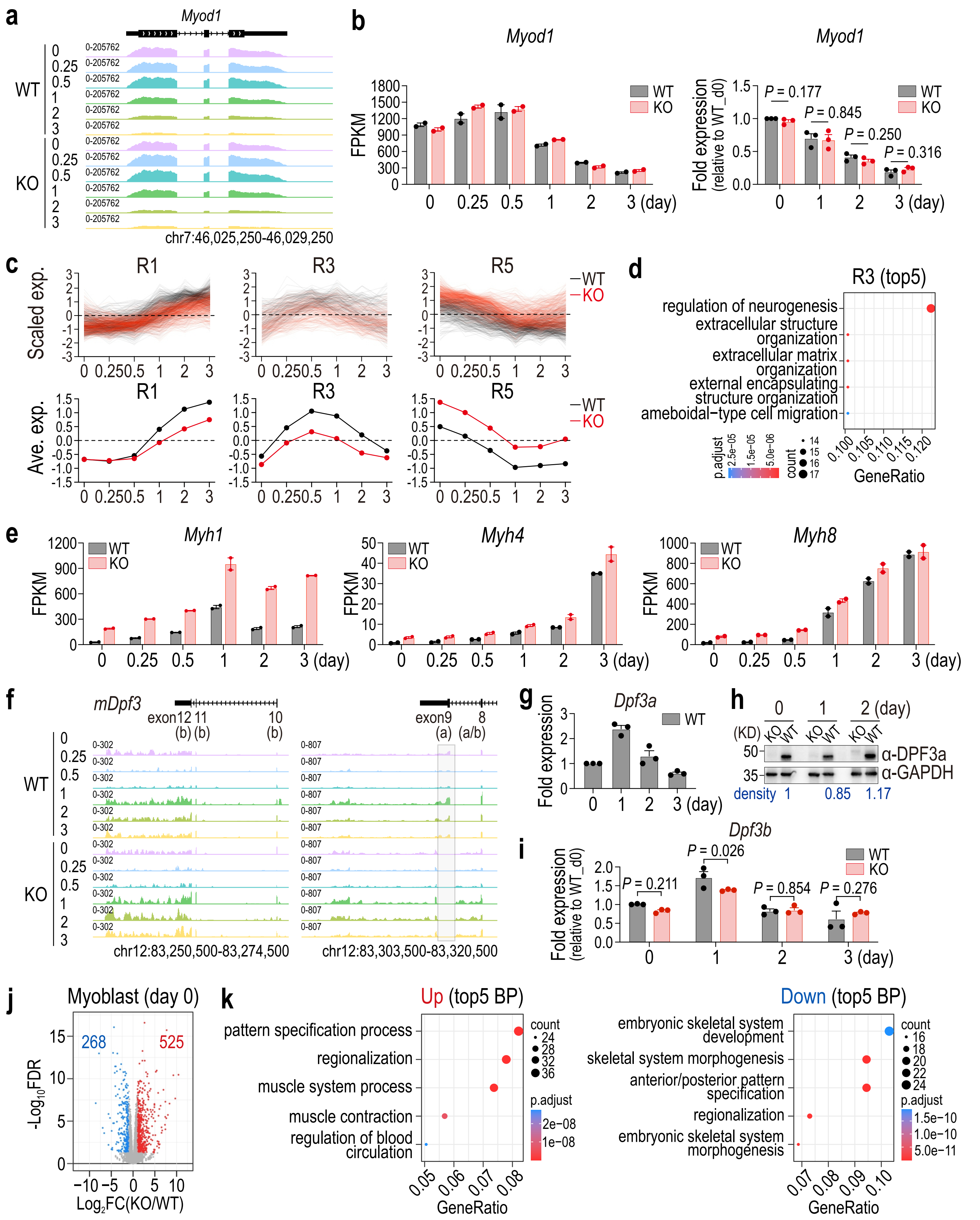

a
Myod1
WT
KO
0
0.25
0.5
1
2
3
0-205762
chr7:46,025,250-46,029,250
b
Myod1
FPKM
WT
KO
3 (day)
Fold expression (relative to WT_d0)
P = 0.177
P = 0.845
P = 0.250
P = 0.316
c
R1
R3
R5
Scaled exp.
Ave. exp.
—WT
—KO
d
R3 (top5)
regulation of neurogenesis
extracellular structure organization
extracellular matrix organization
external encapsulating structure organization
ameboidal−type cell migration
p.adjust
count
GeneRatio
e
Myh1
Myh4
Myh8
FPKM
WT
KO
3 (day)
f
mDpf3
exon12 11 10
(b) (b) (b)
exon9 8
(a) (a/b)
0-302
0-807
chr12:83,250,500-83,274,500
chr12:83,303,500-83,320,500
g
Dpf3a
Fold expression
WT
3 (day)
h
0 1 2 (day)
(KD) KO WT KO WT KO WT
50
35
α-DPF3a
α-GAPDH
density 1 0.85 1.17
i
Dpf3b
Fold expression (relative to WT_d0)
P = 0.211
P = 0.026
P = 0.854
P = 0.276
WT
KO
j
Myoblast (day 0)
268
525
-Log10FDR
Log2FC(KO/WT)
k
Up (top5 BP)
pattern specification process
regionalization
muscle system process
muscle contraction
regulation of blood circulation
count
p.adjust
GeneRatio
Down (top5 BP)
embryonic skeletal system development
skeletal system morphogenesis
anterior/posterior pattern specification
regionalization
embryonic skeletal system morphogenesis

### Figure S3

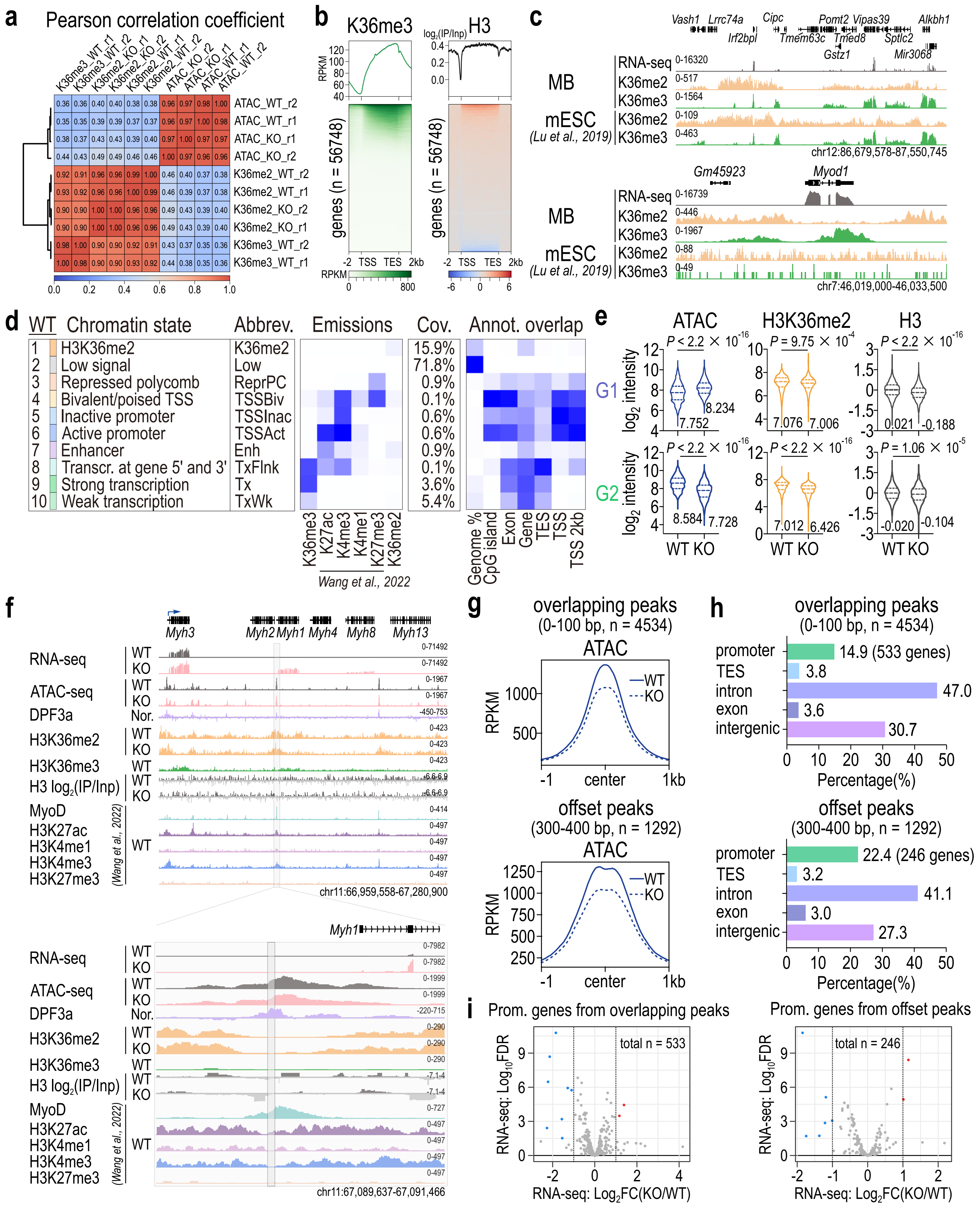

a
Pearson correlation coefficient
K36me3_WT_r1
K36me3_WT_r2
K36me2_KO_r1
K36me2_KO_r2
K36me2_WT_r1
K36me2_WT_r2
ATAC_KO_r2
ATAC_KO_r1
ATAC_WT_r1
ATAC_WT_r2
ATAC_WT_r2
ATAC_WT_r1
ATAC_KO_r1
ATAC_KO_r2
K36me2_WT_r2
K36me2_WT_r1
K36me2_KO_r2
K36me2_KO_r1
K36me3_WT_r2
K36me3_WT_r1
0.0 0.2 0.4 0.6 0.8 1.0
b
K36me3
H3
RPKM
log2(IP/Inp)
genes (n = 56748)
-2 TSS TES 2kb
c
MB
mESC
(Lu et al., 2019)
RNA-seq
K36me2
K36me3
chr12:86,679,578-87,550,745
chr7:46,019,000-46,033,500
Gm45923
Myod1
d
WT Chromatin state
Abbrev.
Emissions
Cov.
Annot. overlap
1 H3K36me2 K36me2 15.9%
2 Low signal Low 71.8%
3 Repressed polycomb ReprPC 0.9%
4 Bivalent/poised TSS TSSBiv 0.1%
5 Inactive promoter TSSInac 0.6%
6 Active promoter TSSAct 0.6%
7 Enhancer Enh 0.9%
8 Transcr. at gene 5' and 3' TxFlnk 0.1%
9 Strong transcription Tx 3.6%
10 Weak transcription TxWk 5.4%
K36me3 K27ac K4me3 K4me1 K27me3 K36me2
Wang et al., 2022
Genome % CpG island Exon Gene TES TSS TSS 2kb
e
ATAC
H3K36me2
H3
G1
G2
log2 intensity
WT KO
f
Myh3 Myh2 Myh1 Myh4 Myh8 Myh13
RNA-seq
ATAC-seq
DPF3a
H3K36me2
H3K36me3
H3 log2(IP/Inp)
MyoD
H3K27ac
H3K4me1
H3K4me3
H3K27me3
(Wang et al., 2022)
chr11:66,959,558-67,280,900
chr11:67,089,637-67,091,466
g
overlapping peaks
(0-100 bp, n = 4534)
ATAC
offset peaks
(300-400 bp, n = 1292)
RPKM
WT
KO
-1 center 1kb
h
promoter 14.9 (533 genes)
TES 3.8
intron 47.0
exon 3.6
intergenic 30.7
promoter 22.4 (246 genes)
TES 3.2
intron 41.1
exon 3.0
intergenic 27.3
Percentage(%)
i
Prom. genes from overlapping peaks
Prom. genes from offset peaks
total n = 533
total n = 246
RNA-seq: Log10FDR
RNA-seq: Log2FC(KO/WT)

### Figure S4

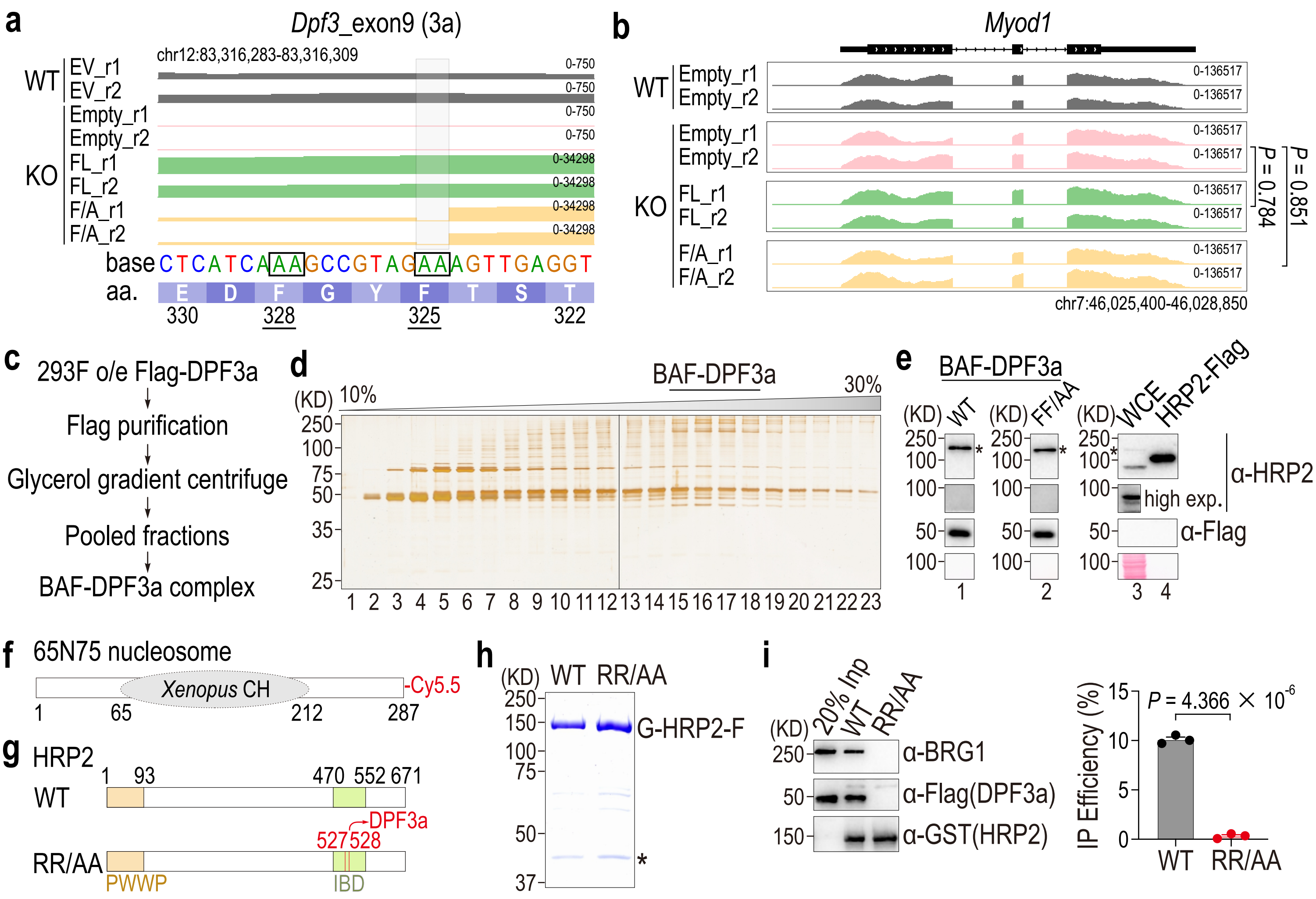

a
Dpf3_exon9 (3a)
chr12:83,316,283-83,316,309
WT
EV_r1
EV_r2
KO
Empty_r1
Empty_r2
FL_r1
FL_r2
F/A_r1
F/A_r2
0-750
0-34298
base CTCATCAAAGCCGTAGAAAGTTGAGGT
aa. E D F G Y F T S T
330 328 325 322
b
Myod1
WT
Empty_r1
Empty_r2
KO
Empty_r1
Empty_r2
FL_r1
FL_r2
F/A_r1
F/A_r2
0-136517
P = 0.784
P = 0.851
chr7:46,025,400-46,028,850
c
293F o/e Flag-DPF3a
Flag purification
Glycerol gradient centrifuge
Pooled fractions
BAF-DPF3a complex
d
BAF-DPF3a
(KD) 10%
30%
250
100
75
50
35
25
1 2 3 4 5 6 7 8 9 10 11 12 13 14 15 16 17 18 19 20 21 22 23
e
BAF-DPF3a
(KD) WT
(KD) FF/AA
(KD) WCE HRP2-Flag
250
100
50
high exp.
α-HRP2
α-Flag
1 2 3 4
f
65N75 nucleosome
Xenopus CH
-Cy5.5
1 65 212 287
g
HRP2
WT
1 93 470 552 671
RR/AA
527528
DPF3a
PWWP
IBD
h
(KD) WT RR/AA
250
150
100
75
50
37
G-HRP2-F
*
i
(KD) 20% Inp WT RR/AA
250
50
150
α-BRG1
α-Flag(DPF3a)
α-GST(HRP2)
IP Efficiency (%)
15
10
5
0
P = 4.366 × 10-6
WT RR/AA

### Figure S5

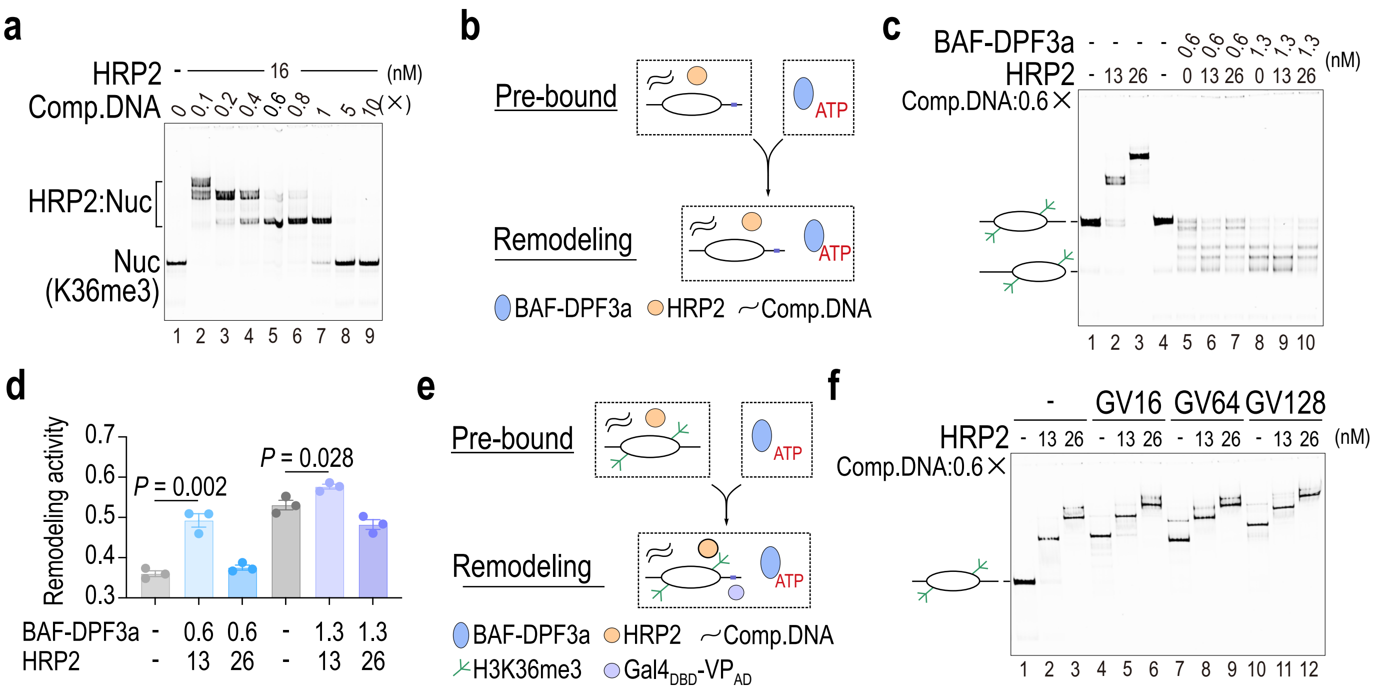
a
HRP2
16
(nM)
Comp.DNA
0 0.1 0.2 0.4 0.6 0.8 1 5 10 (×)
HRP2:Nuc
Nuc
(K36me3)
1 2 3 4 5 6 7 8 9
b
Pre-bound
ATP
Remodeling
ATP
BAF-DPF3a
HRP2
Comp.DNA
c
BAF-DPF3a
0.6 0.6 0.6 1.3 1.3 1.3
HRP2
13 26 0 13 26 0 13 26
(nM)
Comp.DNA:0.6×
1 2 3 4 5 6 7 8 9 10
d
Remodeling activity
0.7
0.6
0.5
0.4
0.3
P = 0.002
P = 0.028
BAF-DPF3a
0.6 0.6 1.3 1.3
HRP2
13 26 13 26
e
Pre-bound
ATP
Remodeling
ATP
BAF-DPF3a
HRP2
Comp.DNA
H3K36me3
Gal4DBD-VPAD
f
GV16
GV64
GV128
HRP2
13 26 13 26 13 26 13 26
(nM)
Comp.DNA:0.6×
1 2 3 4 5 6 7 8 9 10 11 12

### Figure S6

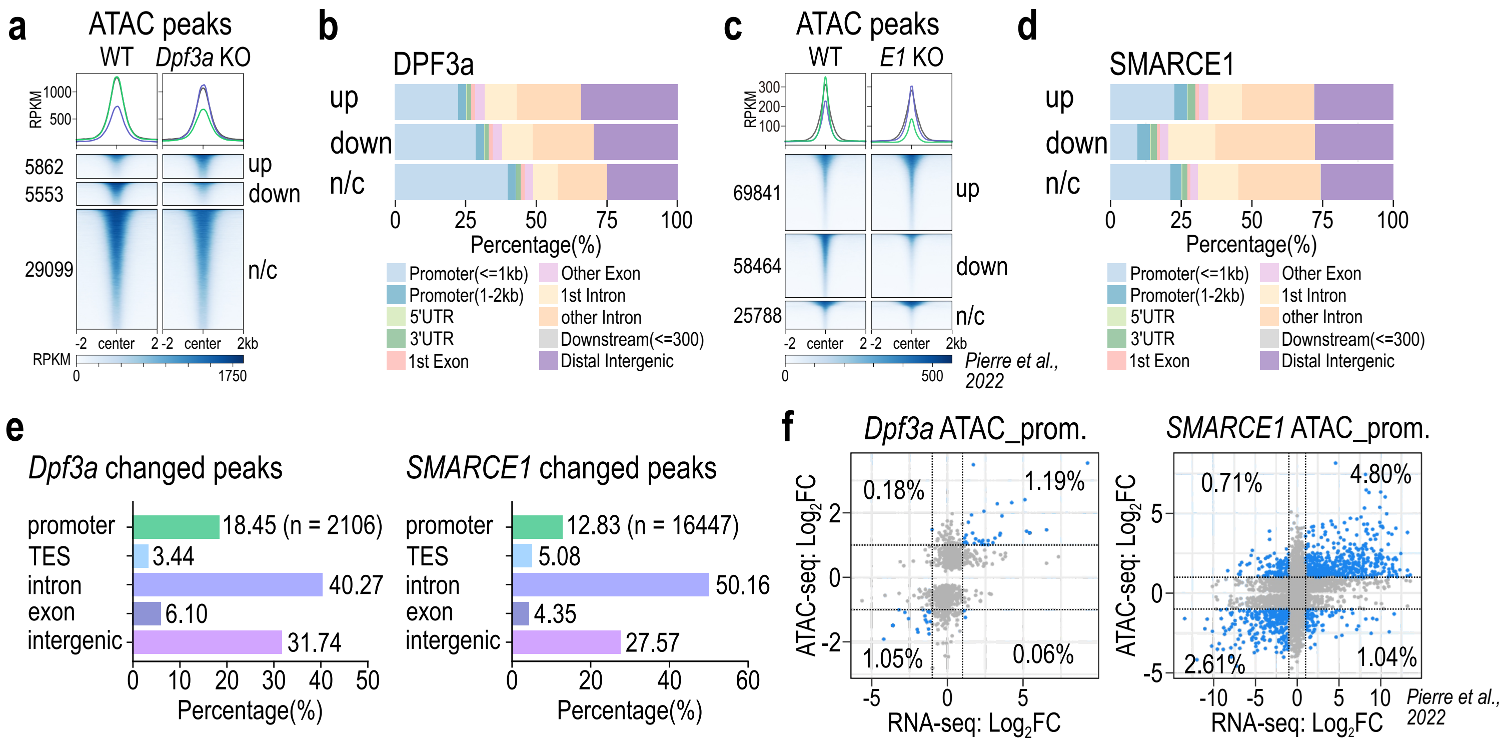


**g**

ATAC_up peaks (DPF3a)

| TF | P value | % of targets | Motif |
|---|---|---|---|
| AP-1 | 1e-636 | 29.77% | TGA TCA |
| RUNX | 1e-62 | 13.01% | ACCACA |
| Myf5 | 1e-55 | 13.37% | CAGCTG |
| MyoD | 1e-52 | 13.98% | CAGCTG |
| MyoG | 1e-49 | 18.08% | CAGCTG |
| CTCF | 1e-11 | 2.20% | TGCC CT TGG |

ATAC_down peaks (DPF3a)

| TF | P value | % of targets | Motif |
|---|---|---|---|
| AP-1 | 1e-169 | 15.81% | TGA TCA |
| Myf5 | 1e-108 | 17.11% | CAGCTG |
| MyoD | 1e-91 | 17.58% | CAGCTG |
| MyoG | 1e-84 | 22.30% | CAGCTG |
| RUNX | 1e-60 | 12.35% | ACCACA |
| CTCF | 1e-37 | 3.48% | TGCC CT TGG |

### Figure S7

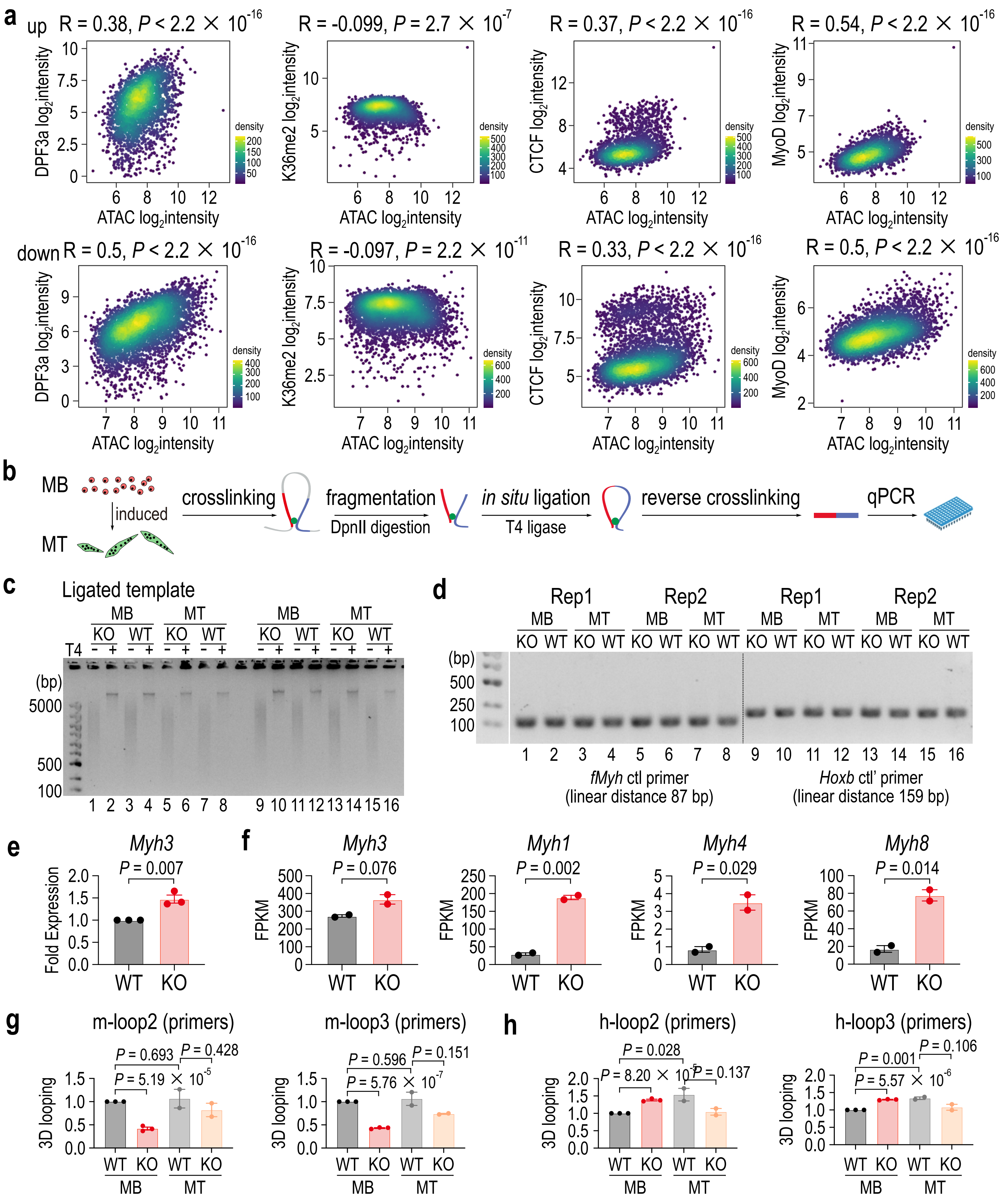

a
up R = 0.38, P < 2.2 × 10^-16
R = -0.099, P = 2.7 × 10^-7
R = 0.37, P < 2.2 × 10^-16
R = 0.54, P < 2.2 × 10^-16
down R = 0.5, P < 2.2 × 10^-16
R = -0.097, P = 2.2 × 10^-11
R = 0.33, P < 2.2 × 10^-16
R = 0.5, P < 2.2 × 10^-16
DPF3a log2intensity
K36me2 log2intensity
CTCF log2intensity
MyoD log2intensity
ATAC log2intensity
density
b
MB
induced
MT
crosslinking
fragmentation
DpnII digestion
in situ ligation
T4 ligase
reverse crosslinking
qPCR
c
Ligated template
MB MT MB MT
KO WT KO WT KO WT KO WT
T4
(bp)
5000
500
100
d
Rep1 Rep2 Rep1 Rep2
(bp)
500
250
100
fMyh ctl primer
(linear distance 87 bp)
Hoxb ctl' primer
(linear distance 159 bp)
e
Myh3
P = 0.007
Fold Expression
WT KO
f
Myh3
P = 0.076
Myh1
P = 0.002
Myh4
P = 0.029
Myh8
P = 0.014
FPKM
g
m-loop2 (primers)
P = 0.693
P = 0.428
P = 5.19 × 10^-5
m-loop3 (primers)
P = 0.596
P = 0.151
P = 5.76 × 10^-7
3D looping
h
h-loop2 (primers)
P = 0.028
P = 8.20 × 10^-5
P = 0.137
h-loop3 (primers)
P = 0.001
P = 0.106
P = 5.57 × 10^-6
WT KO WT KO
MB MT

#### Supplementary Tables

##### Table S1: Resources

| REAGENT or RESOURCE | SOURCE | IDENTIFIER |
|---|---|---|
| **Antibodies** | | |
| Biotin Anti-mouse VCAM-1 | Biolegend | 105704 |
| PE/Cyanine7 Streptavidin | Biolegend | 405206 |
| APC anti-mouse CD31 | Biolegend | 102510 |
| APC anti-mouse CD45 | Biolegend | 103112 |
| FITC Ly-6A/E (Sca-1) (D7) | eBioscience | 11-5981-85 |
| DPF3a | Yupeng Chen Lab | - |
| H3K36me2 | Abcam | ab9049 |
| H3K36me3 | Abcam | ab9050 |
| H3 | Abcam | ab1791-100 |
| CTCF (D31H2) | Cell Signaling Technology | 3418S |
| Anti-Myosin Heavy Chain | Sigma-Aldrich | 05-716-I-100UL |
| HRP2 | ABclonal | A7484 |
| GAPDH | ABclonal | AC002 |
| FLAG-HRP | Sigma | A8592 |
| Mouse IgG-HRP | GE | NA931 |
| Rabbit IgG-HRP | GE | NA934 |
| **Chemicals, peptides, and recombinant proteins** | | |
| Glutathione Sepharose 4B | GE | 17-0756-01 |
| Anti-DYKDDDDK agarose beads | Smart-Lifesciences | SA042100 |
| L-Glutathione Reduced | Sigma | G4251 |
| TRIzol Reagent | Invitrogen | 15596018CN |
| PrimeScript RT reagent Kit | Takara | RR037A |
| Paraformaldehyde | Sigma | 16005 |
| 2×SYBR Green qPCR Master Mix | EZBioscience | A0001-R2 |
| Cardiotoxin | Latoxan | L8102 |
| OCT | Thermo | H-LD-803 |
| Propidium Iodide | Sigma | P4170-10MG |
| ProLong Gold Antifade Mountant | Invitrogen | P36931 |
| Ham's F-10 | Gibco | 11550043 |
| Trypsin-EDTA | Gibco | 25200056 |
| Penicillin-Streptomycin | Gibco | 15140122 |
| Horse serum | Gibco | 16050122 |
| CsCl | Sigma | 289329 |
| Dispase II | Roche | 4942078001 |

| REAGENT or RESOURCE | SOURCE | IDENTIFIER |
|---|---|---|
| Collagenase D | Roche | 11088866001 |
| Recombinant Human FGF | R&D | 233-FB-025 |
| **Critical commercial assays** | | |
| QIAquick PCR Purification Kit | QIAGEN | 28106 |
| QIAquick Gel Extraction Kit | QIAGEN | 28706 |
| RNeasy Mini Kit | QIAGEN | 74104 |
| Hyperactive CUT&Tag Assay Kit | vazyme | TD903 |
| TruePrep DNA Library Prep Kit | vazyme | TD501 |
| VAHTS DNA Library Prep Kit | vazyme | ND607 |
| Stranded mRNA-seq Lib Prep kit | ABclonal | RK20301 |
| **Deposited data** | | |
| ATAC-seq data | This study | CRA037139 |
| RNA-seq data | This study | CRA037139 |
| CUT&Tag data | This study | CRA037139 |
| ChIP-seq data | This study | CRA037139 |
| **Experimental model: Cell lines** | | |
| HEK293A | Ping Hu | RRID:CVCL_6910 |
| HEK293F | Qing Zhong | RRID:CVCL_6642 |
| Recombinant DNA | | |
| DPF3a constructs | This study | N/A |
| HRP2 constructs | This study | N/A |
| $Gal4_{DBD}$ fusion constructs | This study | N/A |
| Software and algorithms | | |
| Rstudio | posit | RRID:SCR_000432 |
| R v.4.4.1 | The R Foundation | RRID:SCR_001905 |
| edgeR v4.4.2 | PMID:19910308 | RRID:SCR_012802 |
| STAR v.2.7.10a | PMID:23104886 | RRID:SCR_004463 |
| samtools v.1.16.1 | PMID:19505943 | RRID:SCR_002105 |
| subread v.2.0.1 | PMID:24227677 | RRID:SCR_012919 |
| deeptools v.3.5.1 | PMID:27079975 | RRID:SCR_ 016366 |
| Bowtie2 v.2.2.5 | PMID:22388286 | RRID:SCR_016368 |
| MACS3 v.3.0.1 | PMID:18798982 | RRID:SCR_013291 |
| Bedtools v.2.30.0 | PMID:20110278 | RRID:SCR_006646 |
| HOMER v.4.11 | PMID:20513432 | RRID:SCR_010881 |

#### Table S2: Plasmids

| Plasmids | Backbone | Plasmid description | Source |
|---|---|---|---|
| pYM118 | pRET | pRET-GST-hHRP2-2Flag | This study |
| pYM146 | pRET | pRET-GST-hHRP2-RR/AA-2Flag | This study |
| pYM237 | pRET | pRET-GST-Gal4$_{DBD}$-G4S-VP16-2Flag | This study |
| pYM238 | pRET | pRET-GST-Gal4$_{DBD}$-G4S-VP64-2Flag | This study |
| pYM239 | pRET | pRET-GST-Gal4$_{DBD}$-G4S-VP128-2Flag | This study |
| pYM101 | pCDH | pCDH-N-3Flag-hDPF3a-puro | This study |
| pYM225 | pCDH | pCDH-N-3Flag-hDPF3aFF/AA -puro | This study |
| pYM296 | pAdTrack | pAdTrack-EGFP | Ping Hu |
| pYM301 | pAdTrack | pAdTrack-N-Flag-mDPF3a-EGFP | This study |
| pYM302 | pAdTrack | pAdTrack-N-Flag-mDPF3aFF/AA -EGFP | This study |
| pYM304 | pAdTrack | pAdRecombinant-Empty-EGFP | This study |
| pYM307 | pAdTrack | pAdRecombinant-N-Flag-mDPF3a-EGFP | This study |
| pYM308 | pAdTrack | pAdRecombinant-N-Flag-mDPF3FF/AA -EGFP | This study |
| pYM136 | pBS | pBL842-di-1x-70bp-linker-gal4 | This study |

#### Table S3: Primers

| Primers | Sequence |
|---|---|
| 5438-DPF3a-KO-F1 | catggaagggaccatcactatgg |
| 5440-DPF3a-KO-F2 | ttgctgggtttccctcactttc |
| 5441-DPF3a-KO-R | gtgagacagttcagccaggacatac |
| 3622-mus-Dpf3a-qPCR-F | cagacgggacagtcattcctaat |
| 3623-mus-Dpf3a-qPCR-R | ctcccaaatgagcagagcgt |
| 3655-mus-Dpf3b-qPCR-F | catgactgaggcagttaagacc |
| 3656-mus-Dpf3b-qPCR-R | cagctcccagcataaatggca |
| 3955-mus-Myod-qPCR-F | atgatgacccgtgtttcgact |
| 3956-mus-Myod-qPCR-R | caccgcagtagggaagtgt |
| 7022-mus-Myh3-qPCR-F | ccaaaacctactgctttgtggt |
| 7023-mus-Myh3-qPCR-R | gggtgggttcatggcataca |
| 3959-mus-Myh1-qPCR-F | cggagtcaggtgaatactcacg |
| 3960-mus-Myh1-qPCR-R | gagcatgagctaaggcactct |
| 9036-mus-Myh8-qPCR-F | ggattgaggcccaaaataagcc |
| 9037-mus-Myh8-qPCR-R | ccatccttgctttgtataacgct |
| 9038-mus-Hoxb2-qPCR-F | accaagaaacccagccaatcc |
| 9039-mus-Hoxb2-qPCR-R | atctgatggtgatccgacctc |
| 9042-mus-Hoxb7-qPCR-F | aagttcggttttcgctccagg |
| 9043-mus-Hoxb7-qPCR-R | acaccccggagaggttctg |

| Primers | Sequence |
|---|---|
| 9044-mus-Hoxb8-qPCR-F | cctgcgccccaattattatga |
| 9045-mus-Hoxb8-qPCR-R | aactcctggatttgcgaaggg |
| 9046-mus-Hoxb9-qPCR-F | tctgggacgcttagcagctat |
| 9047-mus-Hoxb9-qPCR-R | gcccgaaggaaacttggct |
| 8768-3C-fMyh-ctl-F | acaggcgtgaaacctgtgaa |
| 8769-3C-fMyh-ctl-R | gagctaccgaacaaacccca |
| 8925-3C-Hoxb-ctl-F | tcagagacagggtgtagatttcc |
| 8926-3C-Hoxb-ctl-R | gctccttaagagaatccgagtg |
| 8762-3C-fMyh-F | gggtgaccccaaccataaaattg |
| 8767-3C-fMyh-F | aacgggtcttccacgtaacc |
| 8763-3C-fMyh-R | ctccggaggtgcttgatgg |
| 8770-3C-fMyh-R | cctgggtattggtggactataatgag |
| 8775-3C-fMyh-R | ccacctttttgagacatggtctg |
| 8911-3C-Hoxb-F | gccagacccagcttagtgag |
| 8919-3C-Hoxb-F | ctgggtcctcagaggagtg |
| 8914-3C-Hoxb-R | caagtggctgatgggtagtg |
| 8916-3C-Hoxb-R | actctaccattaagctgcacc |
| 5338-Primer-842-mono-F-3 | ccgcgtatagggtccatcaca |
| Primer-Gal4-universal-R-3 | 5'-Cy5.5-agggaacaaaagctggtacccg |
